# Mutanome-guided immunopeptidomics of blood plasma for neoepitope detection in solid tumors is constrained by cfDNA variant calling sensitivity and MS detection limits

**DOI:** 10.64898/2026.09.21.753080

**Authors:** Andreas Kienzle, Jonas P. Becker, Birgit Fendl, Erwin Tomasich, Gerwin Heller, Kathrin Wellach, Kiana Samimi, Britta Eiz-Vesper, Angelika M. Starzer, Julia M. Berger, Vincent Sunder-Plassmann, Barbara Kiesel, Christoph Bock, Georg Widhalm, Matthias Preusser, Anna S. Berghoff, Angelika B. Riemer

## Abstract

**Introduction:** Neoepitopes form the basis of tumor-specific immune responses. Tissue biopsy, the primary source for neoepitope detection, is limited and invasive. Therefore, we aimed to identify neoepitopes by mutanome-guided immunopeptidomics from plasma of cancer patients.

**Methods:** Mass spectrometry (MS) data analysis of HLA ligands from plasma (n = 4) was guided by patient-specific mutanomes of cell-free DNA (cfDNA) from plasma or tumor genomic DNA (tgDNA) from tissue. Matched tumor tissue and healthy donor plasma served as controls. Neoepitopes were validated with synthetic peptides, and immunogenicity was assessed using IFN-γ ELISpot and intracellular cytokine staining.

**Results:** Wild-type immunopeptidomes from tissue and plasma overlapped by 58%, with 91% of plasma HLA ligands rediscovered in tissue. 13 out of 15 tumor-associated HLA ligands detected in plasma were rediscovered in the matching tissue. However, no neoepitopes in plasma were identified by immunopeptidomics guided by cfDNA mutanomes, likely reflecting the limited overlap between cfDNA and tgDNA mutanomes (15%). Using the tgDNA mutanome as a complementary reference, two neoepitopes were detected in one patient’s plasma, albeit at the MS detection limit. Both neoepitopes were also discovered in tissue, along with three tissue-exclusive neoepitopes. Two tissue-exclusive neoepitopes induced antigen-specific T cell responses in healthy donor PBMCs.

**Conclusion:** In summary, plasma immunopeptidomics enables profiling of HLA ligands from wild-type proteins, including TAAs. In principle, neoepitope detection from plasma at the peptide level is feasible, but tissue remains the gold standard for variant calling and neoepitope identification. Improved detection methods may enable minimally invasive approaches in the future.

## Introduction

Tumor antigens, especially mutation-derived neoantigens, are the basis of recognition and subsequent killing of tumor cells by the immune system^1^. Neoantigens mainly arise from non-synonymous somatic mutations and can generate HLA-presented peptides that are recognized by T cells as neoepitopes^2^. The repertoire of all HLA-associated peptides (HLA ligands), including both neoepitopes and endogenous ligands from wild-type proteins, is termed the immunopeptidome^2^.

There are two main strategies for neoepitope discovery: *in silico* prediction of potential neoepitope candidates from tumor sequencing data or direct detection of neoepitopes by MS (immunopeptidomics). For neoepitope prediction, mutations are typically identified by next-generation sequencing of tumor genomic DNA (tgDNA), optionally by integrating RNA sequencing data, prior to HLA-binding prediction and prioritization^3^. However, *in silico* predictions of neoepitopes have a high false-positive rate due to imperfect algorithms^2,4,5^. Mutanome-guided immunopeptidomics, on the other hand, is a strategy to identify actually HLA-presented neoepitopes^2,6^. Here, HLA ligands, including both wild-type epitopes and neoepitopes, are immunoaffinity-purified and analyzed by MS. In parallel, tumor DNA and/or RNA are sequenced to obtain patient-specific mutanomes, which guide the MS-based identification of neoepitopes derived from mutations that are usually absent in standard databases^7^. Candidates are then experimentally validated by comparing a detected hit to a synthetic reference peptide^6^. The immunogenicity of validated neoepitopes is typically evaluated by assessing cytokine secretion (e.g. via ELISpot) or tumor cell killing, reflecting their capacity to elicit an effective anti-tumor immune response^8^.

Low availability of MS-compatible fresh-frozen tumor tissue (as opposed to routinely archived FFPE) poses a significant challenge for neoepitope detection in clinical practice, making liquid biopsy a promising minimally invasive and less restricted alternative^3,9,10^. Direct identification of neoepitopes from plasma via immunopeptidomics has not yet been achieved, although HLA ligands from wild-type proteins, including tumor-associated antigens (TAA), can be detected in complex with soluble HLA from plasma^11–17^.

Blood-derived cell-free DNA (cfDNA), which contains a small fraction of circulating tumor DNA (ctDNA), could provide the mutanome required to guide immunopeptidomics for neoepitope identification in plasma^18–20^. Although the abundance of actual tumor-derived DNA is low and the half-life of cfDNA is short, obtaining the mutanome from liquid biopsy provides a minimally invasive source of mutation information that reflects tumor heterogeneity^18,21^.

The main objective of this study was to identify neoepitopes directly from plasma by immunopeptidomics guided by patient-specific mutanomes from cfDNA. Matched tumor tissue was used to validate both the mutational landscape and the HLA ligands identified from plasma.

## Results

### Pipeline overview for the identification and characterization of neoepitopes from plasma

To explore whether mutation-derived neoepitopes can be directly identified from plasma, we established a mutanome-guided immunopeptidomics workflow that integrates MS-based HLA ligand identification and patient-specific mutanome profiling. Figure 1 provides an overview of the neoepitope identification and characterization pipeline (panel A) and summarizes the analyzed patients (panel B). The cohort included four patients with small cell lung cancer, colorectal cancer, breast cancer, or melanoma, of whom one had received immune checkpoint inhibitor therapy at the time of brain metastasis resection.

**Figure 1:**
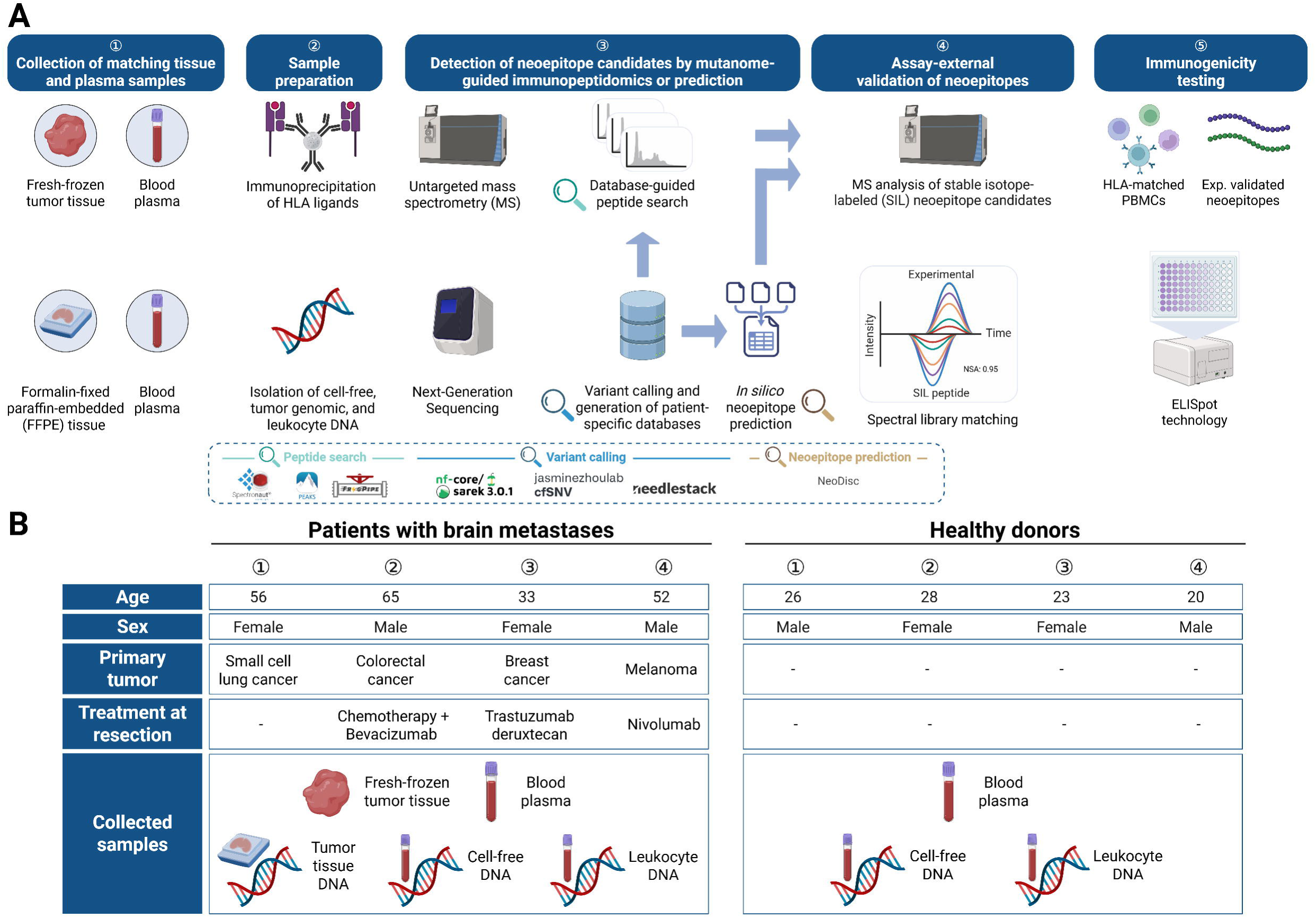
Study overview. **A** Overall pipeline for the identification, validation, and characterization of mutation-derived neoepitopes by mutanome-guided immunopeptidomics. Immunoprecipitated HLA ligands from matched plasma and tumor tissue, as well as from plasma of healthy donors, were analyzed by MS. No tissue was collected from healthy individuals. In parallel, cfDNA, tgDNA, and leukocyte DNA were isolated and subjected to whole-exome sequencing (WES) and variant calling to generate patient-specific mutanomes. Sarek, based on GATK best practices and using Mutect2 as the variant caller, was selected for both cfDNA and tgDNA analysis. For cfDNA samples exclusively, two cfDNA-optimized variant callers (cfSNV, Needlestack) were used. Tailored peptide reference databases, generated from mutanomes, were then used for peptide database search (PEAKS, Spectronaut, and FragPipe) in the MS data. In addition, NeoDisc (variant callers: HaplotypeCaller, Mutect1, Mutect2, and VarScan2) was used for both neoepitope prediction and MS-based immunopeptidomics (FragPipe). Returned mutated peptides were filtered based on HLA-binding predictions (netMHCpan4.1; EL rank ≤ 2%), database query against the HLA Ligand Atlas, and *in silico* spectral library screening (Oktoberfest). Finally, neoepitope candidates were experimentally validated using assay-external synthetic peptides. Immunogenicity was assessed using ELISpot assays. **B** Patient (n = 4) and healthy donor (n = 4) characteristics, primary tumor types, treatment status at the time of brain metastasis resection, and collected sample types used for neoepitope identification and characterization are summarized.

In brief, HLA ligands isolated from plasma were identified using MS-based immunopeptidomics, and the data analysis was guided by tailored peptide reference databases from patient-specific cfDNA mutanomes. Detected neoepitope candidates were validated by comparison to external synthetic peptide reference libraries, and their immunogenicity was assessed. Matched tumor tissues from brain metastatic lesions were included to validate the plasma-derived immunopeptidomes and mutanomes, while plasma from healthy donors (n = 4) served as a negative control.

### High rediscovery rate of wild-type and tumor-associated HLA ligands from plasma in matching tumor tissue

To validate plasma as a source for HLA ligand discovery, we first assessed the overlap in the wild-type immunopeptidomes between matched plasma and metastatic tissue samples of four cancer patients using data-independent acquisition (DIA-)MS immunopeptidomics. Additionally, we analyzed plasma from four healthy individuals to assess possible disease- or treatment-associated differences.

On average, 6,780 (range: 3,339-10,525) wild-type HLA ligands were identified in the plasma of cancer patients, 9,871 (range: 6,987-14,224) in the matching tissue, and 4,988 (range: 4,450-6,228) in plasma samples from healthy individuals (Figure 2 A). There was no significant difference between the number of wild-type HLA ligands identified in the plasma and matching tissue of patients, or between plasma from patients and plasma from healthy individuals (t-test, p > 0.05).

**Figure 2:**
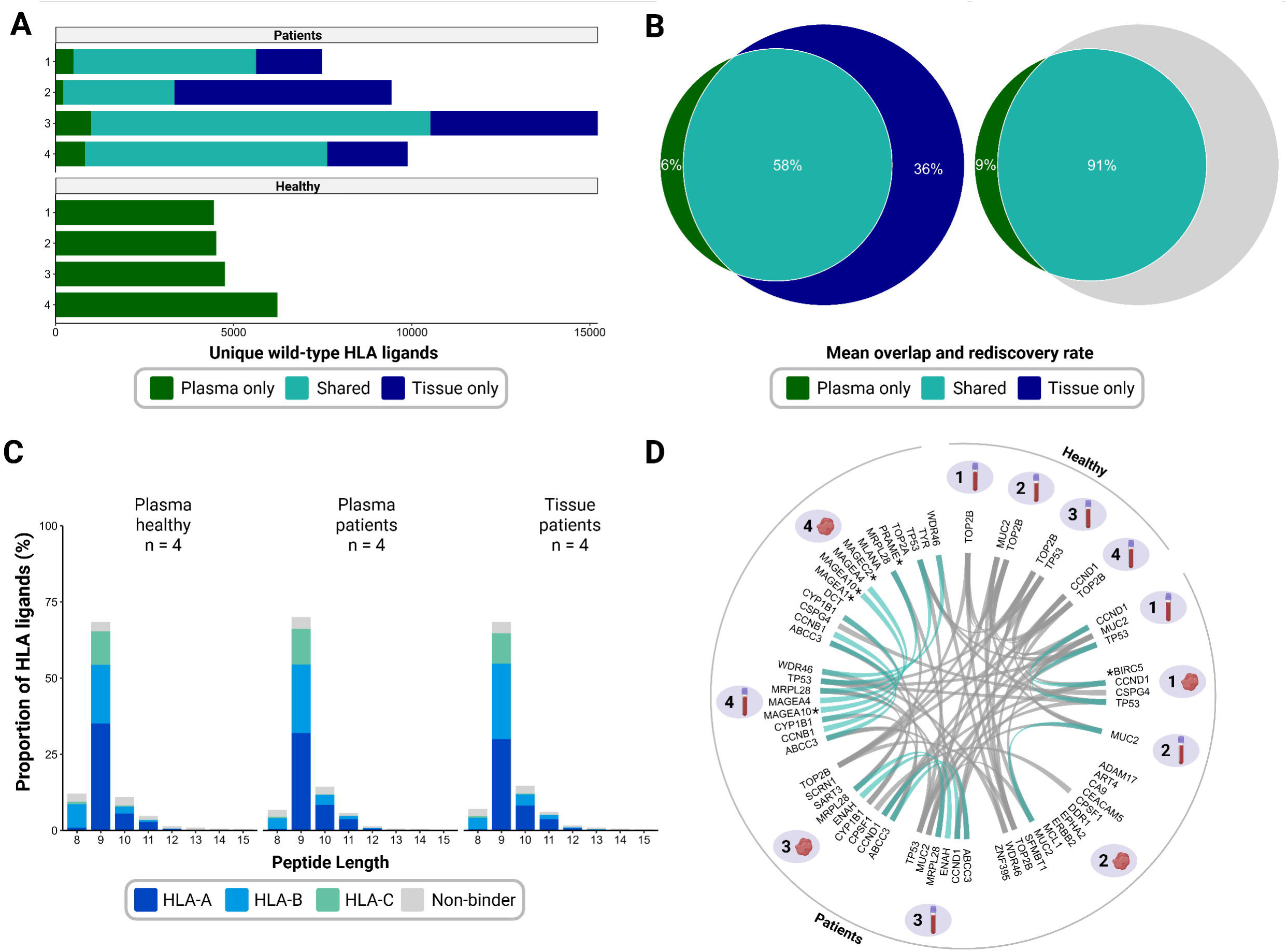
Immunopeptidomics of tissue and plasma for the discovery of HLA ligands from wild-type proteins and tumor-associated antigens. **A** Direct identification of wild-type HLA ligands from plasma and/or tumor tissue of patients (n = 4) and healthy individuals (n = 4) via MS-based immunopeptidomics using the reviewed UniProtKB/Swiss-Prot human reference proteome (20,387 entries). HLA ligands detectable only in plasma are colored green, HLA ligands detected both in a patient’s tumor tissue and plasma are colored turquoise, and HLA ligands only detected in tumor tissue are colored blue. **B** Patient-averaged proportions (%) of wild-type HLA ligands exclusively detected in plasma (green), exclusively detected in matched tissue (blue), or shared between both (turquoise). The left panel shows the overall overlap of HLA ligands between plasma and tissue, whereas the right panel shows the proportion of HLA ligands from plasma rediscovered in matched tissue. **C** Summarized length distribution and relative proportions of best-fit HLA alleles (predicted by NetMHCpan4.1) of wild-type HLA ligands from immunopeptidomic analysis of tumor tissue and/or plasma of patients and healthy individuals. Peptides with an EL rank > 2% were classified as predicted non-binders. **D** Circos plot of source genes encoding tumor-associated antigen (TAA)-derived HLA ligands in plasma and matched tissue. Turquoise links indicate TAAs shared between plasma and tissue from the same patient, whereas grey links indicate TAAs shared across different patients. Asterisks mark cancer testis antigens as defined by Shraibman et al.^17^.

The patient-averaged overlap of identical wild-type HLA ligands between tissue and plasma (colored turquoise in Figure 2 A and Figure 2 B left panel) was 58% (ranging from 33% to 69%). 91% (ranging from 89% to 93%) of wild-type HLA ligands from plasma were rediscovered in the matching tissue of the same individuals, indicating that the wild-type immunopeptidome from plasma largely reflects that of the tissue (Figure 2 B right panel).

Next, we compared the sequence and binding features of the wild-type immunopeptidomes between tissue and plasma. Tissue- and plasma-derived wild-type HLA ligands of patients and healthy individuals featured a typical HLA ligand length distribution of mainly (69±13%) nonamers (Figure 2 C, Supplementary Figure 1 A). The majority of peptides were predicted binders (netMHCpan 4.1) for HLA-A (45±1%), followed by HLA-B (32±2%), and HLA-C with the fewest binders (12±1%), which reflects the expected distribution. The proportion of predicted non-binders was not significantly different between the plasma of patients (11±1%), the plasma of healthy individuals (13±1%), and the tissue of patients (12±1%; Wilcox test: p > 0.05). Using sequence clustering (*GibbsCluster 2.0*), we observed highly similar but not completely identical HLA-binding motifs in the immunopeptidomes from plasma and tissue (Supplementary Figure 2). Two patients exhibited more clusters in plasma, one patient showed equal numbers of clusters in plasma and tissue, and one patient had more clusters in tissue. In two patients, clusters of exclusively HLA-C-predicted binders were found in plasma but not in matched tissue. For example, wild-type peptides from the plasma of Patient 1 were segregated into four clusters corresponding to the HLA-binding motifs of HLA-A*02:05, HLA-A*11:01, HLA-B*13:02, HLA-B*39:01, HLA-C*06:02, and HLA-C*12:03, which are the HLA alleles expressed by this patient (Supplementary Table 1). Of the four clusters identified in the plasma immunopeptidome of Patient 1, three were also present in the tissue immunopeptidome, whereas one cluster (for HLA-C*06:02 and HLA-C*12:03) appeared exclusively in plasma.

Lastly, as an *in silico* quality control, we used the deep learning peptide retention time (RT) predictor *DeepLC* to validate the MS data of wild-type HLA ligands^28^. Overall, we observed a high correlation (ρ ≥ 0.94, p ≤ 0.001) of predicted and experimentally measured RTs for both tissue- and plasma-derived wild-type HLA ligands, further supporting the quality of our immunopeptidomics data (Supplementary Figure 1 B).

To support the premise that tumor-derived peptides can be detected in plasma, the presence of HLA ligands from TAAs was investigated (Figure 2 D, Supplementary Table 2).

Among the patients’ plasma and tumor immunopeptidomes, 42 distinct HLA ligands originating from 34 known TAAs were identified. 13 out of 15 distinct HLA ligands from plasma were also rediscovered in the matching tissue. All eleven distinct TAAs with HLA ligand support from plasma (except MUC2 and/or TP53 in Patient 1 and 3) were rediscovered in the matching tissue (turquoise links). Notably, one of the TAAs (MAGE-A10) from plasma was a cancer testis antigen (CTA), a class of TAA characterized by germline-restricted expression and aberrant overexpression in tumors (asterisks in Figure 2 D highlight CTAs)^17^. Seven distinct HLA ligands from four TAAs (MUC2, TP53, TOP2B, and CCND1) were detected in the plasma of healthy individuals, of which three HLA ligands and all four TAAs were also detected in tissue and plasma samples from patients. Detection of TAAs in healthy individuals is not surprising, because TAAs are by definition not tumor-restricted and can show low expression in healthy tissues^22^.

In sum, plasma and tissue immunopeptidomes were highly similar, with comparable yields, high rediscovery rates of plasma-detected wild-type HLA ligands in tissue, and typical peptide length distributions, with only minor differences in predicted HLA-binding motifs. Importantly, HLA ligands originating from TAAs were also detectable in plasma, which were largely rediscovered in the matching tumor tissue, supporting the potential of plasma immunopeptidomics as a source of tumor-derived epitopes.

### Limited mutational concordance between cfDNA and tumor tissue DNA

The immunopeptidomics- and prediction-based discovery of mutation-derived neoepitopes requires tailored peptide reference databases derived from non-synonymous variants, which are typically patient-specific. Encouraged by the substantial overlap of the plasma and tissue immunopeptidomes and to establish a fully blood-based workflow without relying on tissue biopsies, mutanomes were obtained from cfDNA using matched leukocyte DNA as a reference. In this study, non-synonymous variants (i.e. missense mutations, insertions, deletions) were identified using standard variant calling workflows implemented in NeoDisc (HaplotypeCaller, VarScan2, Mutect1, Mutect2 with custom hard filtering) and Sarek (Mutect2), as well as cfDNA-optimized variant callers (Needlestack, cfSNV). To determine whether the mutanome detected in cfDNA reflects that of the tumor, we compared the overlap and yield of non-synonymous variants between cfDNA and tgDNA, the standard material for mutational profiling.

First, we analyzed the default outputs of each variant calling pipeline, retaining only variants passing the caller-specific built-in quality filters. On average, more non-synonymous variants were detected in cfDNA (mean = 839, range: 778-906) compared to tgDNA across all four patients (n = 751, range: 545-1091) (Figure 3 A). The mutanome overlap between cfDNA and tgDNA was limited, with only 15% of variants on average per patient being shared (turquoise in Figure 3 A). Notably, a total of 490 and 294 variants detected in cfDNA and tgDNA of cancer patients, respectively, were also detected in cfDNA of healthy individuals (turquoise in Figure 3 B). In addition, most non-shared cfDNA variants were exclusively detected by NeoDisc or Needlestack, raising the question of whether this is due to the orthogonal nature of the variant calling algorithms or the true hit validity (Figure 3 C).

**Figure 3:**
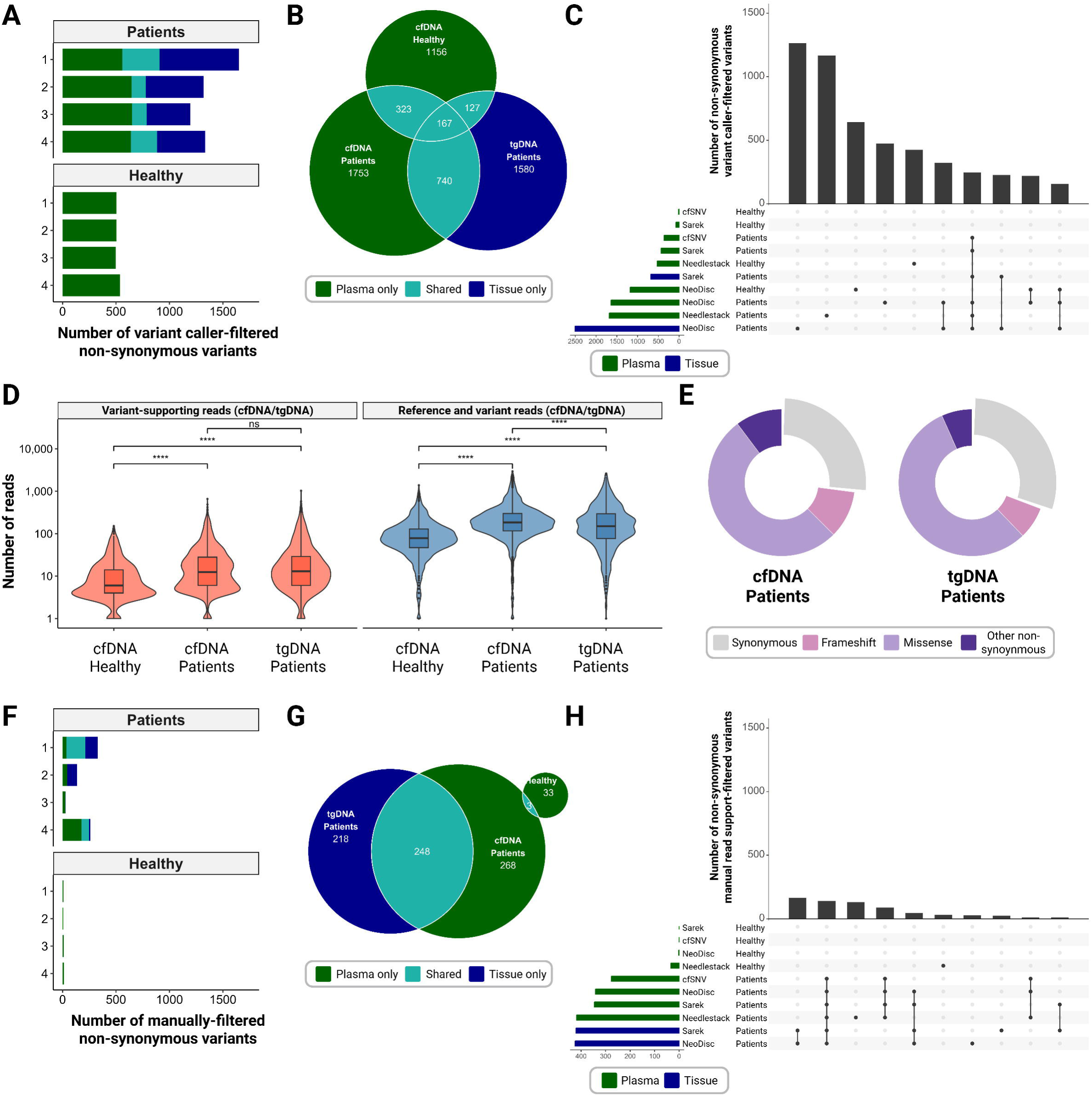
Variant calling from cfDNA (plasma) and tgDNA (tissue) **A** Overlap and yield of non-synonymous variants from tissue (tumor genomic DNA, tgDNA) and/or plasma (cell-free DNA, cfDNA) of cancer patients (n = 4) and healthy individuals (n = 4) passing the caller-specific built-in quality filters. Variants detected only in cfDNA are shown in green, variants detected in both cfDNA and tgDNA in turquoise, and variants detected only in tgDNA in blue. Variant calling for healthy individuals was performed only from cfDNA, as no tissue was collected. **B** Overlap of total non-synonymous variants, passing the caller-specific built-in quality filters, between cfDNA and tgDNA of patients and healthy individuals. Overlaps are shown in turquoise. **C** Top 10 intersections of non-synonymous variants, passing the caller-specific built-in quality filters, across cfDNA (green) and tgDNA (blue) samples from cancer patients and healthy individuals detected by four variant calling pipelines (cfSNV, Needlestack, NeoDisc, Sarek). cfSNV and Needlestack were exclusively used for cfDNA samples. **D** Evaluation of variant read support (left) and total read depth at the variant locus (right) for variants passing the caller-specific built-in quality filters from cfDNA and tgDNA. Box plots show medians and interquartile ranges of the read support. P-values from Wilcox tests with Bonferroni correction are indicated above brackets (**** = p < 0.0001; ns = not significant). **E** Distribution of synonymous and non-synonymous mutations detected in cancer patients (n = 4) after caller-specific built-in quality filtering. **F** Overlap and yield of non-synonymous variants from tissue (tgDNA) and/or plasma (cfDNA) of cancer patients (n = 4) and healthy individuals (n = 4), which were manually filtered based on variant read support (>10 reads on tumor DNA and 0 on leukocyte DNA), total read depth at the variant locus (>15 reads), and on StrandOddsRatio, SOR (<4 for single-nucleotide variants and <10 for insertions and deletions). Variants exclusive to cfDNA are shown in green, shared variants in turquoise, and variants exclusive to tgDNA in blue. **G** Overlap of total non-synonymous variants, passing the manual filters, between cfDNA and tgDNA of patients and/or healthy individuals. Overlaps are shown in turquoise. **H** Top 10 interactions of non-synonymous variants passing manual filters (read coverage, SOR) across cfDNA (green) and tgDNA (blue) samples from cancer patients and healthy donors, detected by four pipelines (cfSNV, Needlestack, NeoDisc, Sarek). cfSNV and Needlestack were used only for cfDNA analysis.

We therefore also examined the variant read support and the total read depth (including both variant and reference reads) at the variant loci in cfDNA and tgDNA of patients and healthy individuals. In the case of cfDNA, patients showed on average higher variant read support (median: 13±40) and higher total read depth (184±285) at the variant position compared to healthy individuals (median: 6±17 and 78±119, respectively; Figure 3 D; both p < 0.0001 from Wilcox test with Bonferroni correction). This aligned with the significantly higher cfDNA concentrations in patients (median: 13 ng/mL, range: 6-34 ng/mL plasma) compared to healthy individuals (median: 4 ng/mL, range: 2-6 ng/mL plasma; Wilcox test, p = 0.03), likely reflecting the presence of tumor-derived cfDNA. The variant read support was comparable between cfDNA and tgDNA of patients (Wilcox test with Bonferroni correction, p > 0.05). Further, the distribution of variant types between cfDNA and tgDNA in all patients was highly concordant with mainly missense (55±2%) and synonymous mutations (29±2%) (Figure 3 E). About 9±1% of mutations constituted frameshift variants. The proportion of other non-synonymous mutations (e.g. in-frame insertions and deletions or nonsense variants) was 8±1%. Using mutational signature analysis (Supplementary Figure 3 A-B) and base change profiling (Supplementary Figure 3 C) of synonymous and non-synonymous variants, we observed comparable signatures across cfDNA and tgDNA samples and only low fractions of sequencing artifact signatures, such as AR1 signatures or formalin-induced C>T changes in tgDNA. Signature AC5 (COSMIC SBS5), suggested to represent age-dependent spontaneous mutagenesis in tumor and in healthy somatic cells, was most dominant across all samples^23^. Notably, cfDNA and tgDNA samples from melanoma Patient 4 exhibited both a prominent AC7 signature (COSMIC SBS7), which is associated with ultraviolet-related cutaneous melanoma^24^.

To address low-confidence variants and increase specificity, variants were filtered based on their locus-specific read support in both tumor and matched normal DNA, and strand bias filtering (StrandOddsRatio, SOR) was applied to remove artifacts and sequencing errors. In addition, variants reported by NeoDisc were retained only if supported by at least two variant callers (consensus calling), as HaplotypeCaller is not optimized for low-frequency somatic variant detection^25^. On the one hand, this improved the patient-averaged overlap of variants between cfDNA and tgDNA from 15% to 20% and also reduced the number of variants detected in cfDNA of healthy individuals from a total of 1,773 to 36 variants (Figure 3 F and G), which were primarily detected by Needlestack (Figure 3 H). Also, a high proportion of manually filtered variants were co-detected by multiple variant calling pipelines (Figure 3 H). On the other hand, variant overlaps between cfDNA and tgDNA were observed in only two out of four patients. Notably, Patient 1 showed 54% variant overlap between cfDNA and tgDNA, with an 83% rediscovery rate of cfDNA variants in matching tgDNA. In addition, the number of variants detected in the cfDNA of patients was reduced from 2,983 to 519 variants.

In summary, the mutanome detected in cfDNA only partially reflected that of the tumor tissue DNA. Stringent filtering improved specificity but substantially reduced variant yield and did not consistently increase concordance across patients. To preserve sensitivity, we retained all variants that passed caller-specific default filters and performed additional locus-specific read support assessment for neoepitope-underlying variants from downstream immunopeptidomic analyses.

### Neoepitope discovery via mutanome-guided immunopeptidomics in matched plasma and tissue samples

#### cfDNA-guided immunopeptidomics did not result in neoepitope detection in blood plasma

Variants passing the caller-specific quality filters across variant calling pipelines from cfDNA of four cancer patients and four healthy individuals were then used to build tailored databases to identify potential neoepitopes in the DIA-MS data of the respective plasma sample.

NeoDisc was used for both library-free MS-based neoepitope detection (FragPipe) and *in silico* neoepitope prediction, whereas outputs from Sarek, Needlestack, and cfSNV were used for MS-guided neoepitope detection (Spectronaut, PEAKS, FragPipe) in library-free or library-guided modes (based on wild-type experimental and neoepitope *in silico* libraries). All MS-based searches used the reviewed UniProtKB/Swiss-Prot human reference proteome as the canonical background database, in line with best practice in immunopeptidomics. Unlike MS-based search engines (Spectronaut, PEAKS, FragPipe), NeoDisc for *in silico* neoepitope prediction operates solely on DNA input, yielding candidates without a priori assignment to a sample source (blood/plasma). Retention times were therefore predicted using DeepLC, and candidates were subsequently evaluated in both plasma and tissue DIA-MS data.

The library-free strategy returned eleven distinct mutated peptides in patients and none in healthy individuals. The library-guided strategy yielded 201 distinct mutated peptides in patients and 106 in healthy individuals. The prediction strategy returned 392 distinct mutated peptides in patients and 379 in healthy individuals. All peptides were evaluated *in silico* based on the Normalized Spectral Angle (NSA), by measuring the similarity between the experimentally observed fragment spectrum for a candidate in the DIA-MS data and a reference library spectrum predicted by *Oktoberfest* (*in silico* NSA ≥ 0.7; transitions ≥ 3). Only five neoepitope candidates, MS-reported in the plasma and by using reference databases from cfDNA, passed the *in silico* evaluation, without any candidates being returned from healthy individuals (green dots in the top part of Figure 4 A).

**Figure 4:**
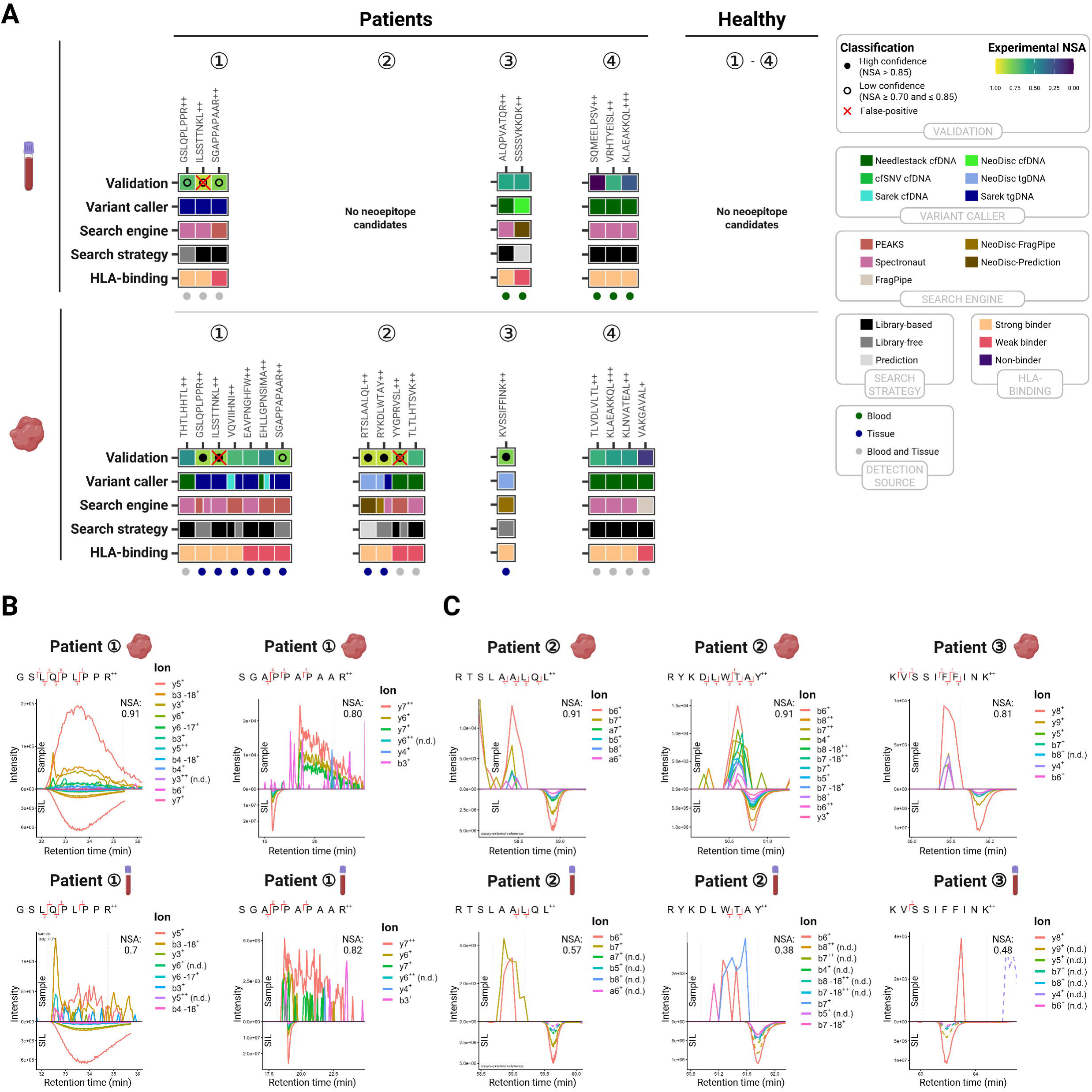
Identification and experimental validation of neoepitope candidates from tissue and plasma. **A** Neoepitope candidates returned from mutanome-guided immunopeptidomics of plasma and/or tissue samples from cancer patients. All candidates in DIA-MS data were experimentally evaluated by matching to spectral libraries generated from SIL peptides. The normalized spectral angle (NSA) scores the similarity of the detected peptide’s fragmentation pattern to the reference library, where 1 indicates perfect identity and 0 total dissimilarity. Validated detections are indicated by a full (NSA > 0.85; full dot) or hollow (NSA ≥ 0.70 and ≤ 0.85) dot. Candidates manually classified as false-positives (e.g. due to RT discrepancies to SIL peptides) were marked with a red cross. Candidates without a hollow or full dot were non-confirmable. For each candidate, the search strategy and engine (both MS- and prediction-based) and the variant calling tool (cfDNA: green shades, tgDNA: blue shades) are specified. Candidates were further classified into weak (EL rank 0.5-2%) and strong binders (EL rank ≤ 0.5%) based on HLA-binding predictions (netMHCpan 4.1). Green dots indicate candidates that were detected in the plasma and by using reference databases from cfDNA, representing detections identifiable solely based on plasma-derived data. Blue dots indicate candidates that are completely identifiable from tissue-based detections. Candidates with grey dots were identifiable by complementary analyses, for instance, peptides detected in plasma whose underlying genetic variants were called from tissue sequencing. **B** Experimental validation of two neoepitope candidates detected in tissue and plasma, shown in panel A, by evaluating the intensity of fragment ions, corresponding to extracted ion chromatograms (XICs), and the retention times. Upper plots show mirrored transitions, used to determine the NSA, of the neoepitopes detected in DIA-MS of tissue against corresponding SIL peptides. Lower plots show the same neoepitopes detected in DIA-MS of matching plasma, also plotted against the spectral library from SIL peptides. **C** Neoepitope candidates identified exclusively in tissue samples using patient-specific tgDNA mutanomes, which were not confidently detected in the matching plasma. Candidates were validated by comparison with SIL peptide spectral libraries.

Finally, *in silico* screened neoepitope candidates were experimentally validated using synthetic peptides. Of note, no targeted PRM-MS analysis, with or without spiked-in stable isotope-labeled (SIL) peptides, was performed due to the lack of remaining patient material. Instead, the NSA of a candidate in the DIA-MS data was obtained by comparing to a reference library spectrum from an assay-external targeted MS analysis of SIL peptides alone (c.f. “Validation” in Figure 4 A). After the experimental validation step, candidates were classified as either non-confirmable (NSA < 0.7; transitions < 3), low confidence (NSA ≥ 0.70 and ≤ 0.85; hollow dot), high confidence (NSA > 0.85; full dot), or false-positive (red cross) hits^26–29^. Strikingly, none of the five neoepitope candidates returned by cfDNA-guided immunopeptidomics could be experimentally validated using SIL peptides (exp. NSA < 0.7 and/or transitions < 3).

#### tgDNA-guided immunopeptidomics enabled the identification of neoepitopes in the blood plasma of one patient

Based on our observation that cfDNA mutanomes only partially reflected those of matching tgDNA, we expanded our approach to include mutanomes from tgDNA to build complementary personalized reference databases to search DIA-MS raw data of plasma samples (grey dots in the top part of Figure 4 A). Three new neoepitope candidates from plasma, which were all detected in Patient 1, passed our previously applied thresholds for the *in silico* and experimental validation (Figure 4 A; Supplementary Table 3): Neoepitope *GSLQPLPPR* (MS search engine: *Spectronaut*; variant calling pipeline: *Sarek*; parental gene: missense mutation in *MMAB*) was a low confidence hit (experimental NSA: 0.7; five unique transitions). A similar RT to that of the SIL peptide was observed, showing a wide retention window from 32 to 36 minutes (Figure 4 B). The same peptide was detected with high confidence (exp. NSA 0.91, seven unique transitions) and the same broad signal in the matching tissue (Figure 4 B; PEAKS and Spectronaut). This neoepitope had a strong predicted HLA-binding affinity to HLA-A*11:01 (EL rank: 0.107%). Neoepitope *SGAPPAPAAR* (*PEAKS*; *Sarek*; frameshift variant in *INSM1*) was a low confidence hit (exp. NSA: 0.82; four unique transitions). This neoepitope was predicted to be a weak HLA-A*11:01 binder (EL rank: 1.747%). This neoepitope was also co-detected in the matching tissue (exp. NSA: 0.8; four unique transitions; PEAKS). In the tissue, the mean RT of this neoepitope differed from that of the SIL peptide (19 vs. 16 min; Figure 4 B). We still considered *SGAPPAPAAR* as a true hit, as we hypothesize that small RT shifts (<3 min) can occur in chromatographic regions exhibiting strong peptide co-elution, which could delay elution of the target peptide relative to the SIL standard^30^, although we used RT calibrants to synchronize RTs between measurements. Indeed, we observed the maximum peptide identifications eluting around 20 minutes in the tissue and plasma samples, which could lead to column saturation effects resulting in broad signals and delayed elution (Supplementary Figure 5). Neoepitope candidate *ILSSTTNKL* (exp. NSA of 0.9, but only three unique transitions) was classified as a false-positive hit as the RT differed from that of the SIL peptide (27 min vs. 33 min; Supplementary Figure 6).

#### Immunopeptidomics of matched tissue confirms all plasma-detected neoepitopes and reveals tissue⍰exclusive neoepitopes

Next, we used the patient-specific reference databases from both cfDNA and tgDNA mutanomes to search neoepitopes in matching metastatic tissue, to potentially detect neoepitopes that were not identified in the immunopeptidomics analysis of plasma (lower part of Figure 4 A). Using this approach, 16 candidates passed the *in silico* evaluation (*in silico* NSA ≥ 0.7; transitions ≥ 3). Candidates completely identifiable in tissue by immunopeptidomics guided by tgDNA databases are annotated by blue dots, while candidates returned from MS analysis of tissue and guided by cfDNA database are annotated by grey dots. Out of these, five neoepitope candidates could be experimentally validated by SIL peptides. The two tissue detections in Patient 1 (*GSLQPLPPR*, *SGAPPAPAAR*) were the same as those from plasma (Figure 4 B and Supplementary Table 3). The detection *ILSSTTNKL* in Patient 1 was again classified as a false-positive hit due to a RT mismatch with the SIL peptide (Supplementary Figure 6). Three new detections, in Patient 2 (*RTSLAALQL, RYKDLWTAY*) and Patient 3 (*KVSSIFFINK*), are shown in Figure 4 C. In the tissue of Patient 2, *RTSLAALQL* was identified by *NeoDisc* (prediction-based) using the patient-specific tgDNA mutanome (HaplotypeCaller, Mutect2, VarScan; frameshift in *ZFP36L2)* and classified as a high confidence hit (exp. NSA: 0.91; four unique transitions). Although this peptide was not confidently detected in the matching plasma sample, two high-ranking transitions from the SIL peptide reference spectrum were observed in the plasma DIA-MS2 spectra, suggesting that the detection may have been just below the detection limit. This neoepitope candidate featured a strong HLA-binding capacity to HLA-C*15:17 (EL rank: 0.108%). Neoepitope *RYKDLWTAY* from Patient 2 was returned by both *NeoDisc* (FragPipe) and *Spectronaut* based on the tgDNA mutanome (HaplotypeCaller, Mutect1, VarScan and/or Mutect2; missense variant from *SUPT6H*) as a high confidence hit (exp. NSA: 0.91; six unique transitions). This neoepitope featured a strong HLA-binding capacity to HLA-C*07:01 (EL rank: 0.161%). The signals for this neoepitope in the matching plasma sample were low (exp. NSA: 0.38; three unique transitions). In addition, neoepitope *YYGPRVSL* was detected as a low confidence hit (exp. NSA: 0.81; five unique transitions) by *PEAKS* and guided by the cfDNA-mutanome from *Needlestack* (insertion in *OR5H1*). However, because the RTs differed from those of the SIL peptide (45 vs. 51 min), this candidate was classified as a false-positive detection (Supplementary Figure 6). In the tissue of Patient 3, another neoepitope (*KVSSIFFINK*) was validated with low confidence (exp. NSA 0.81; six unique transitions), deriving from *NeoDisc* (FragPipe) using the tgDNA mutanome (HaplotypeCaller; frameshift in *TAF1B*). Like for *RTSLAALQL* (Patient 2), *KVSSIFFINK* could not be confidently detected in the matching plasma sample of Patient 3, but transition rank 1 (y8) was observed at similar RTs as the SIL peptide (Figure 4 C). This neoepitope featured a strong HLA-binding capacity to HLA-A*03:01 (EL rank: 0.079%).

Notably, not all variants resulting in MS-confirmed neoepitopes would have passed the manual filters based on read coverage and strand bias for the tumor and leukocyte DNA, and consensus calling (NeoDisc only). The variant underlying *GSLQPLPPR* (Sarek: GRCh38 chr12:109567094 C T) would have been excluded, because less than ten reads supported the variant (Supplementary Table 3, Supplementary Figure 4). The variant underlying *KVSSIFFINK*, reported by NeoDisc (HaplotypeCaller), was not supported by consensus calling and less than ten reads supported the variant (hg19 chr2: 9989570 TA T). Further, two reads supporting the variant in the matching leukocyte DNA, used as the reference, were observed.

In summary, cfDNA-guided immunopeptidomics did not enable the detection of neoepitopes from plasma or tissue in our dataset. tgDNA-guided immunopeptidomics allowed the detection of two mutation-derived neoepitopes (*GSLQPLPPR*, *SGAPPAPAAR*) directly from plasma, but at the MS detection limit. The same neoepitopes (*GSLQPLPPR*, *SGAPPAPAAR*) and three tissue-exclusive neoepitopes (*RYKDLWTAY, RTSLAALQL, KVSSIFFINK*) were confidently detected in tissue samples from three out of four patients using patient-specific tgDNA-derived mutanomes.

### Detection of antigen-specific T cell responses to a subset of validated neoepitopes

Lastly, we evaluated the *in vitro* immunogenicity of the identified neoepitopes to determine whether they can potentially trigger antigen-specific T cell responses^31^. To this end, expanded PBMCs from healthy donors (n = 7) were stimulated with synthetic peptides and analyzed by IFN-γ ELISpot and intracellular cytokine staining (ICS) assays. Of note, PBMCs from healthy donors were used because no PBMCs from the respective cancer patients were available. In addition, T cell exhaustion and low T cell frequency were often observed in patient-derived PBMCs, which would limit the sensitivity of response detection. For these reasons, the use of HLA-matched healthy donor PBMCs represents an established approach for assessing the general immunogenicity of neoepitopes^32–35^. Donors were selected based on HLA matching (exact allele or supertype) with the predicted restricting alleles (Supplementary Table 1, Supplementary Table 4), with HLA supertypes defining groups of alleles that bind and present overlapping peptide repertoires^36^.

Stimulation with *RTSLAALQL* (SI = 8.12, SFU = 964 per 10□PBMCs) and *KVSSIFFINK* (SI = 30.22, SFU = 3,220 per 10□ PBMCs) induced peptide-specific IFN-γ secretion in ELISpot (SI > 3 and number of SFU per 10□ PBMCs > 200) in one of the seven donors (Figure 5 A and B). Epitope-specific CD8⁺ T cell activation was confirmed by IFN-γ and TNF-α production after peptide stimulation in ICS (Figure 5 C). *RTSLAALQL* and *KVSSIFFINK* stimulation resulted in 9.53% and 7.16% double-positive (IFN-γ⁺ TNF-α⁺) cells compared to DMSO background controls (0.59% and 0.48%), respectively. *GSLQPLPPR* and *SGAPPAPAAR*, which were both detected in plasma, did not induce IFN-γ secretion in the PBMCs of the tested healthy donor. *RYKDLWTAY*, which was exclusively detected in tissue, similarly failed to show detectable immunogenicity. Overall, two out of five neoepitopes detected in tissue induced measurable antigen-specific T cell responses.

**Figure 5:**
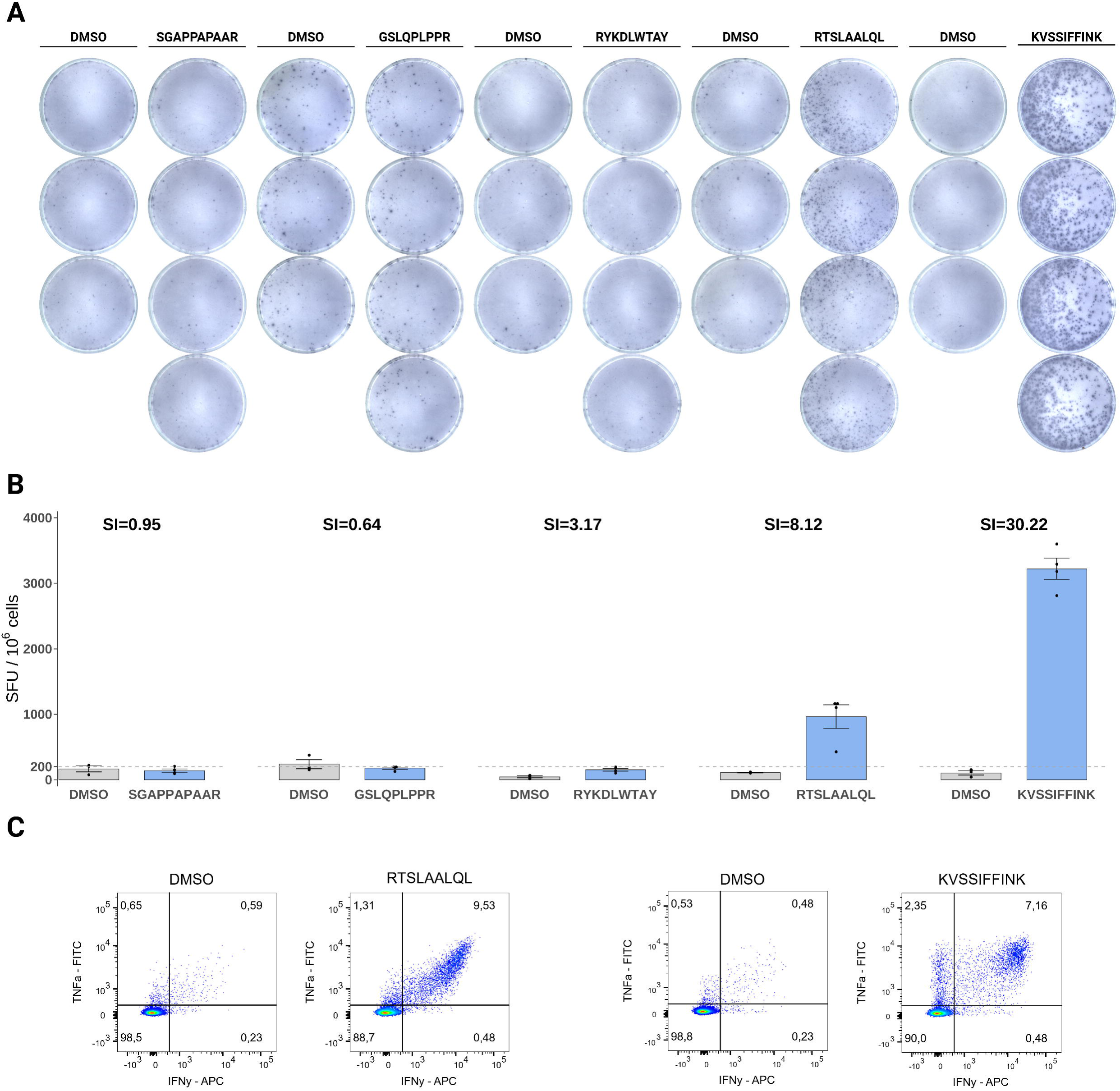
Immunogenicity assessment of neoepitopes detected by immunopeptidomics. **A** IFN-γ ELISpot results from *in vitro* expanded PBMCs of an HLA-(supertype-)matched healthy donor stimulated with immunopeptidomics-detected peptides or DMSO (negative control). The HLA-genotype of this healthy individual was HLA-A*11:01, HLA-A*23:01, HLA-B*18:03, HLA-B*44:03, HLA-C*04:01, HLA-C*07:01. **B** Quantification of IFN-γ spot-forming units (SFU) per 10 PBMCs (mean ± SD, n = 4 for peptide treatment or n = 3 for DMSO) from panel A. SI refers to the stimulation index. Induction of IFN-γ secretion was classified as positive if both the SI exceeded 3 and the number of SFU per 10 PBMCs was greater than 200 (SFU threshold indicated by the dashed line). **C** Intracellular cytokine staining for IFN-γ and TNF-α in CD8⁺ T cells upon peptide or DMSO stimulation. The percentages of IFN-γ⁺, TNF-α⁺, and double-positive cells are shown in the respective plot quadrant.

## Discussion

Neoepitopes hold potential as therapeutic targets and predictive biomarkers in immunotherapy, but their detection in cancer patients is challenging due to limited tissue availability. In this study, mutanome-guided immunopeptidomics was used for minimally invasive neoepitope detection, which allowed the identification of two neoepitopes in the plasma of one patient that were also detectable in matching tissue. However, high confidence MS identification was not achieved, and a reference database from tissue DNA rather than cfDNA was required to guide immunopeptidomics, making it only partly plasma-based.

This was consistent with our finding that immunopeptidomes from tissue and plasma were highly similar, whereas cfDNA and tgDNA mutanomes showed only partial concordance. In the case of the immunopeptidomes, wild-type HLA ligands overlapped substantially between tissue and plasma and exhibited the typical HLA ligand features, aligning well with previously published data on wild-type HLA ligands from plasma^17,37^. Slight differences in HLA-binding motif clustering between plasma and tissue may indicate that plasma represents a global immunopeptidome from multiple tissue sources, supporting liquid biopsy as a more comprehensive alternative to single-site tissue biopsies^11^. For example, in the plasma of two patients, additional sequence clusters for HLA-C were observed that were absent in the matching tissue. As altered HLA-C expression has been implicated in tumor immune evasion, these clusters could also reflect tumor-driven changes in HLA-C peptide presentation, although other non-tumor sources cannot be excluded^38^. In line with Shraibman et al.^17^, the plasma immunopeptidome contained HLA ligands from several TAAs, which were also present in the matching tumor tissue. In addition, HLA peptides derived from the highly immunogenic CTA MAGE-A10 were identified, supporting the premise that clinically relevant tumor-derived peptides can be captured in plasma^39^. While wild-type and tumor-associated HLA ligands were largely shared between tissue and plasma, mutanomes obtained from cfDNA samples only partially reflected those of the matching tgDNA samples. This aligns with previous studies using WES on matched tissue DNA and cfDNA samples, which reported mutanome concordances as low as 20%^40,41^. In addition, variants were also detected in cfDNA of healthy controls, underscoring the difficulty of distinguishing artifacts from true mutations, even when using matched leukocyte DNA as a reference. Though seemingly artefactual, at least one non-synonymous mutation has been reported in 60% of cfDNA from healthy individuals, which mainly originate (>80%) from hematopoietic cells with high turnover rates^42,43^. Other studies even report up to seven mutations per Mb in non-cancer individuals^44^.

Likely reflecting the low concordance between cfDNA and tgDNA mutanomes and the presence of variants lacking tumor specificity, no neoepitopes could be reliably identified using immunopeptidomics guided by mutanomes from cfDNA. We further speculate that the high dilution factor of HLA ligands in circulation, the relatively limited plasma volume used for analysis, and the potential loss of weak binders in plasma samples may have additionally limited neoepitope detectability^45^. By using personalized reference databases built from matching tgDNA mutanomes to guide immunopeptidomics, two neoepitopes in the plasma of one patient (*GSLQPLPPR*, *SGAPPAPAAR*) were detected. The co-elution with synthetic peptides and rediscovery of those peptides in the matching tissue support true detections, despite low transition counts and spectral matches to SIL peptides. Of note, *GSLQPLPPR* appears in an alternative isoform of MAP3K15 (Q6ZN16-3), but the annotation derives from a single *in silico* translation of cDNA ORFs without experimental validation (ECO:0000303) and is absent from the reviewed UniProtKB/Swiss-Prot human reference proteome used here for database searching and filtering, consistent with best practice in immunopeptidomics^46^.

The triggers for the release and the immunological functions of soluble HLAs complexed with (mutated) HLA ligands are not fully understood, but tumor cells are thought to actively secrete them to evade immune recognition^47^. The neoepitope *GSLQPLPPR* originated from a missense mutation in *MMAB*, a gene encoding a mitochondrial enzyme critical for vitamin B12 metabolism that is linked to metastatic progression^48,49^. In the same patient with small cell lung carcinoma (SCLC), the neoepitope *SGAPPAPAAR* was detected, which derived from the transcription factor *INSM1*. *INSM1* serves as a diagnostic biomarker with high sensitivity and specificity for neuroendocrine tumors, including SCLC^50,51^.

Nevertheless, tissue currently remains the gold standard for neoepitope identification. In our study, three additional neoepitopes (*RYKDLWTAY*, *RTSLAALQL, KVSSIFFINK*) were exclusively detected in tissue, again guided by mutanomes from tgDNA. The parental genes of the neoepitopes *RYKDLWTAY* (*SUPT6H*, encoding a histone chaperone) and *RTSLAALQL* (*ZFP36L2*, encoding an RNA-binding protein) promote tumor initiation and progression in colorectal cancer, respectively^52,53^. The neoepitope *KVSSIFFINK*, derived from *TAF1B*, has been previously reported in IEDB as a non-canonical immunopeptide of unassigned protein origin. However, its sequence corresponds to a frameshift product of a coding mononucleotide repeat in *TAF1B*, a recurrent mutation hotspot in mismatch-repair-deficient cancers^54^. *KVSSIFFINK* is also contained within the *TAF1B* frameshift peptide evaluated in a clinical MSI-H cancer vaccine trial (NCT01461148)^55^. This supports a mutation-derived rather than a non-mutational, non-canonical origin of *KVSSIFFINK*.

Of note, manual variant filtering to increase specificity based on read support, strand bias, and consensus calling (restricted to NeoDisc) would have reduced our set of five validated neoepitopes to only three candidates (*SGAPPAPAAR*, *RTSLAALQL*, *RYKDLWTAY*). For example, the variant giving rise to *KVSSIFFINK*, reported by NeoDisc, showed low variant read support and additionally lacked consensus calling support, although this does not necessarily indicate a false positive, as limited consensus across variant callers has been reported elsewhere^56,57^. In the matching leukocyte DNA, two variant-supporting reads at low VAF (5%) at this locus were observed, suggesting a sequencing artifact rather than a true germline variant. Despite limited DNA-level evidence, immunopeptidomics provided an orthogonal validation of the mutation at the peptide level for *KVSSIFFINK* and *GSLQPLPPR.* Finally, we tested all five detected neoepitopes for immunogenicity with HLA-matched PBMCs from healthy donors, as evidence of immunogenicity for a given epitope is desirable for designing informed next-generation clinical trials, particularly for neoepitope-based vaccines^32–35^. Two tissue-exclusive neoepitopes (*RTSLAALQL* and *KVSSIFFINK*) induced measurable antigen-specific T cell responses. Although functional immune activation was not observed for plasma-detected neoepitopes, this does not preclude their potential relevance, because ELISpot sensitivity for low-frequency T cells and other *ex vivo* factors may limit detection of immunoreactivity^58^. In addition, reactivity was tested in a small healthy donor panel, with only one donor showing reactivity even for the tissue-derived neoepitopes, suggesting that a larger donor cohort might have revealed additional reactive cases. Of note, this analysis does not assess whether the patients had T cells reactive against these peptides, but just that they are immunogenic in principle.

Other limitations of our study include the small size of this pilot study, the limited confidence in MS detections of plasma-detected neoepitopes, and the unavailability of remaining patient material for targeted MS validation. As plasma was collected after surgery, the level of tumor-derived cfDNA and soluble HLA complexes could have been substantially reduced. Future studies should therefore collect plasma before surgical resection to maximize recovery of tumor-derived cfDNA and soluble HLA-bound peptides.

Taken together, our study demonstrates that mutanome-guided immunopeptidomics for the direct identification of neoepitopes from plasma in cancer patients is, in principle, technically feasible, but to date only when guided by mutanomes from tgDNA. To advance clinical translation, future studies should focus on further methodological refinement, such as implementing next-generation MS platforms and more sensitive variant calling pipelines, as well as the clinical evaluation of plasma-derived neoepitopes in larger, prospectively collected patient cohorts. The integration of immunopeptidomics with liquid biopsy holds promise for minimally invasive and personalized cancer immunotherapy.

## Conclusion

Immunopeptidomics of liquid biopsy enables minimally invasive detection of HLA ligands from TAAs, but neoepitope identification in blood plasma is challenging and currently relies on mutanomes from tumor tissue DNA. Further refinement through the integration of next-generation mass spectrometry and (cfDNA-)optimized variant calling will be essential to establish its utility in personalized cancer immunotherapy.

## Supporting information

Highlights

## Data availability

The mass spectrometry data have been deposited to the ProteomeXchange Consortium via the PRIDE partner repository with the dataset identifier PXD082101^59^.

## Material and methods

### Cohorts and collection of patient material

The project has received ethical approvals from the ethics committee of the *Medical University of Vienna (*1164/2019, 1616/2020, 1832/2021) for all procedures involved after informed consent and in compliance with the Declaration of Helsinki, relevant laws, and institutional guidelines. Brain metastatic lesions of four cancer patients (Figure 1 B) were surgically resected at the *Department of Neurosurgery* of the *Medical University of Vienna / Vienna General Hospital*. Arterial plasma was collected within 24 hours post metastasis resection. The control cohort comprised four healthy volunteers (Figure 1 B). Venous plasma was drawn at a single time point at the *Department of Medicine I (Medical University of Vienna / Vienna General Hospital)*.

### DNA isolation

For cfDNA isolation, plasma from patients (mean volume: 9 ± 2 mL) and healthy volunteers (6 ± 1 mL) was generated by centrifugation of whole blood in *PAXgene Blood DNA tubes* (Qiagen, Germantown, MD) for 20 min at 300xg and re-centrifugation for 10 min at 5,000xg at room temperature (RT). The *QIAamp circulating nucleic acid kit* (Qiagen, Germantown, MD) was used for cfDNA isolation. Tumor DNA from metastatic lesions of patients was isolated from 30 µm shavings of archived FFPE tissue slides according to the *Maxwell® FFPE Plus DNA Kit* (Promega, Madison, WI) and using a *Maxwell® RSC* instrument (Promega, Madison, WI). Leukocyte DNA from patients and healthy volunteers was isolated from 6 mL K3-EDTA blood (Greiner Bio-One, Kremsmünster, Austria) using the *E.Z.N.A.® SQ Blood DNA Kit* (Omega Bio-Tek, Norcross, GA, United States). Isolated DNA was purity-checked by a NanoDrop device (absorbance ratio 1.8-1.9 for 260/280 nm).

### Next-generation sequencing and variant calling

Exome sequencing (37 Mb) for all DNA samples (cfDNA, leukocyte DNA, and tgDNA) was conducted with 100 base pairs paired-end reads on a NovaSeq 6000 (Illumina, San Diego, California, USA). Levels of sequencing artifacts arising from deamination of cytosines in FFPE tissue and oxidation of guanine to 8-oxoguanine were estimated using GATK4 CollectSequencingArtifactMetrics. Pre-adapter summary metrics from all samples (cfDNA, leukocyte DNA, and tumor tissue DNA) yielded Phred-scaled TOTAL_QSCOREs ≥ 50 for OxoG artifacts and 100 for deamination artifacts across all samples, indicating no evidence of FFPE-induced cytosine deamination or oxidative damage artifacts. Variants were called from cfDNA using cfSNV (version 0.99.0; default settings and using recommended sample-specific parameters for VAF and read support based on sample coverage; GRCh38), Needlestack (version 1.1; default settings; GRCh38), Sarek (version 3.5.1; caller: Mutect2; GRCh38), and NeoDisc (version 1.7.0, caller: HaplotypeCaller, Mutect1, Mutect2, and Varscan2; hg19)^60–62^. Sarek and NeoDisc were also used for variant calling from WES of tgDNA. Matching leukocyte DNA was used as a reference in all cases. The NeoDisc variant calling output was converted from hg19 to GRCh38 using *GATK LiftoverVcf* to allow comparison to the other variant callers. Ensembl Variant Effect Predictor VEP (version 109.3) was used for annotation. For an exploratory sub-analysis, variants were also manually filtered based on the variant read support (>10 reads on tumor DNA and 0 on leukocyte DNA), total read depth at the variant locus (>15 reads) in tumor DNA and leukocyte DNA, and on the StrandOddsRatio for the variant (<4 for SNVs and <10 for InDels) in tumor DNA, which was calculated using GATK VariantAnnotator. In addition, NeoDisc variants were further filtered if supported by at least two variant callers (consensus calling).

### Mutational signature analysis

COSMIC mutational signature analysis was performed using the YAPSA R package on single-nucleotide variants (SNVs) generated from all variant callers (either all variants passing the caller-specific built-in quality filters [Supplementary Figure 3 A-C] or those passing the manual filters using read coverage and strand bias filters [Supplementary Figure 3 D-F]). 96-channel mutation catalogs were based on the hg38 reference genome. To account for exome-specific biases, capture-corrected normalization factors (hs37d5) were applied. Signature contributions were estimated by fitting reference COSMIC signatures, including both biological and technical signatures. Normalized signature exposures were compared by hierarchical clustering using Manhattan distance (k = 3).

### HLA-genotyping

HLA-genotyping was conducted at the *Department for Blood Group Serology and Transfusion Medicine* (*General Hospital of Vienna*) as described in Fischer et al. (2019)^63^. In brief, PCR-based long-range amplifications of HLA-A, -B, and -C genes from DNA isolated from K3-EDTA blood were achieved by using GoTaq Long PCR Mastermix together with HLA-loci specific primers (5’ UTR to 3’ UTR). Amplicons with a size of >5000 bp were fragmented with enzymes and barcoded. Fragments of approx. 400 bp size were subsequently selected on an E-Gel electrophoresis system. Clonal amplification was performed by an emulsion PCR of 26 pMol fragments, prior cleaning steps, enrichment, and loading of the ion sphere particles onto a 520 Chip. The sequencing was performed on a S5 sequencing device from Thermo Fisher. Bioinformatic analysis of the reads was performed by using the TypeStream NGS Analysis software (Thermo Fisher) together with the GenDX NGS (GenDX Utrecht) software.

### Immunopeptidomics

Membrane-bound and soluble HLA complexes were immunoprecipitated from fresh-frozen tumor tissue and/or plasma of patients (n = 4) and healthy volunteers (n = 4) using a high-throughput 96-well plate format initially described for the identification of neoepitopes from tissue^37^. No tissue was collected and analyzed from healthy individuals.

In detail, whole K3-EDTA plasma (mean volume: 16 ± 2 mL) from patients and healthy volunteers was centrifuged at 1,200xg for 10 min, supplemented 1:1000 with a protease inhibitor cocktail (Sigma-Aldrich, St. Louis, MO), and again centrifuged at 12,000xg for 10 min at 4°C. Fresh-frozen tissue from patients was homogenized with an *ULTRA-TURRAX homogenizer* (IKA®-Werke GmbH & Co. KG, Darmstadt, Germany) in lysis buffer (25% sodium deoxycholate, 0.2 mM iodoacetamide, 1 mM EDTA, 1:200 Protease Inhibitors Cocktail, 1 mM Phenylmethylsulfonyl fluoride, and 1% octyl-beta-d glucopyranoside in PBS). The lysis mixture was centrifuged at 40,000xg for 30 min at 4°C. Cleared plasma or purified tissue lysate sequentially passed two stacked 96-well micro-plates (Agilent, Böblingen, Germany), which were pre-filled with protein-A sepharose or anti-HLA class I antibody (clone: W6/32) crosslinked protein A-sepharose beads. The immunoprecipitate was sequentially washed with four times wash solution A (150 mM NaCl and 20 mM Tris-HCI Buffer pH 8), wash solution B (400 mM NaCl and 20 mM Tris-HCI Buffer pH 8), and wash solution A as well as twice with wash solution C (400 mM NaCl and 20 mM Tris-HCI Buffer pH 8) using a *Waters Positive Pressure-96 Processor*. Elution into a *Sep-Pak tC_18_ 100 mg Sorbent* 96-well plate (Waters, Milford, MA) was performed with 1% TFA prior desalting with 0.1% TFA and final elution with 28% ACN in 0.1% TFA. Dried samples were resuspended in 5% ACN with 0.1% TFA and 100 fmol/µL *Pierce™ Peptide RT Calibration Mixture* (Thermo Scientific™, Waltham, AS).

All samples were analyzed with a *Thermo Ultimate 3000* coupled to an *Exploris 480* (Orbitrap MS) equipped with a FAIMS Pro interface (Thermo Fisher Scientific) and using a compensation voltage of -50 V or -65 V, as described previously^64^.

### Direct identification of wild-type HLA ligands and mutation-derived neoepitopes

FAIMS-DIA MS data was searched against the reviewed UniProtKB/Swiss-Prot human reference proteome (retrieved: 21.10.2021; 20,387 entries) for the identification of wild-type peptides using Spectronaut (version 17.1; Biognosys). The peptide level false-discovery rate (FDR) was set to 1%, and carbamidomethylation and methionine oxidation were defined as variable modifications. Replicate injections (CV-50V and CV-65V) were grouped into a single condition per sample type (plasma or tissue) and individual. Method evaluation was enabled to allow separate analysis of tissue and plasma samples. Peptides were further analyzed using NetMHCpan-4.1 binding predictions, *GibbsCluster 2.0* clustering of peptide sequences, and RT prediction by DeepLC.

For the discovery of mutation-derived neoepitopes, three different search strategies were applied: First, FAIMS-DIA data was analyzed library-free against patient-specific peptide reference databases in *Spectronaut* (version 17.1), *PEAKS* (version 12), and *FragPipe* (version 20). In-house Python scripts were used to generate immunopeptidomics databases from VEP-annotated variants that passed the caller-specific quality filters (Needlestack, Sarek, and cfSNV), retaining protein-coding missense, and frameshift variants from either tgDNA or cfDNA. Second, custom *in silico* spectral libraries were generated from the same patient-specific databases using *Oktoberfest* (prediction model*: Prosit_2020_intensity_HCD, enzyme: “no_enzyme”, precursor charge: 1-4)*^65^. To reduce the search space, only peptides (8 to 15 amino acids) with an EL rank ≤2.0% (*netMHCpan* version 4.1) were included. *Spectronaut*, *PEAKS*, and *FragPipe* were then used as MS search engines guided by the *in silico* spectral libraries. Experimental libraries derived from wild-type HLA ligands after DIA-MS (*Spectronaut;* reviewed UniProtKB/Swiss-Prot human reference proteome) were included as background. For both the library-free and in silico library-based strategies, search parameters for the Spectronaut search were set to non-enzyme digestion and to 1% FDR at the peptide level. Carbamidomethyl and Methionine oxidation were included as variable modifications. The reviewed UniProtKB/Swiss-Prot human reference proteome (entries: 20,387 entries) was included as a reference. For PEAKS, the enzyme specificity was set to none, precursor peptide mass error tolerances were set to 10 ppm and 0.02 Da for fragment ions. Protein N-term acetylation, methionine oxidation, and carbamidomethylation were set as variable modifications, with three possible modifications allowed per peptide (8-15 amino acids). The peptide FDR threshold was set to 1%. The reviewed UniProtKB/Swiss-Prot human reference proteome (entries: 20,387 entries) and a contamination database (common Repository of Adventitious Proteins [cRAP]) from the Global Proteome Machine (GPM) organization (retrieved: 04.08.2024; 125 entries) were included in the analysis. For FragPipe the default nonspecific-HLA workflow was used with following adaptions: Database searching was run without enzyme specificity and peptide length was set to 8 to 15 amino acids. Precursor ion tolerance and fragment ion tolerance were set to ±10 ppm and 0.2 Da, respectively. Carbamidomethylation was set to a fixed modification. Methionine oxidation and acetylation at protein N terminus were set to variable modifications. The FDR was controlled by Percolator, and identifications were filtered to 1% at the PSM level. Of note, for Spectronaut, PEAKS, and FragPipe searches, run-to-run matching was disabled, while Method Evaluation in Spectronaut was enabled to ensure independent analysis of tissue and plasma samples without cross-run imputation. As a third discovery strategy, NeoDisc (version 1.7.0) was run in sensitive mode to identify neoepitopes from DIA MS raw data using its own variant calling pipeline (Mutect1, Mutect2, VarScan2, HaplotypeCaller). Default parameters were used. The capture kit was set to “Twist_HCEP_V1”.

### Validation of mutation-derived neoepitope candidates

All neoepitope candidates, returned from the three discovery strategies, were queried against the HLA Ligand Atlas^66^ (retrieved: 10.03.2025; 223,246 entries). As a first screening step, neoepitope candidates were *in silico* evaluated in *Skyline* (version 22)^67^ based on their normalized spectral angle match (*in silico* NSA) with predicted spectra from *Oktoberfest* (*in silico* NSA ≥ 0.5 or ≥ 0.7; transitions ≥ 3) as well as a good match of the LC experimental RT with the predicted RTs (deepLC)^68^.

Lastly, identified neoepitopes in DIA-MS data were experimentally validated by spectra comparison with SIL peptides using *Skyline*. SIL peptides, containing two heavy amino acids (lysine +8 Da, arginine +10 Da, isoleucine/leucine +7 Da, valine +6 Da, proline +6 Da, or phenylalanine +10 Da, depending on peptide sequence), were obtained from *JPT Peptide Technologies* (purity >70%). First, the RT for each target peptide, dissolved in 5% ACN and 0.1% TFA and together with peptide RT calibration (PRTC) mixture (Pierce Biotechnology), was determined by LC-MS analysis in DDA mode with an inclusion list of all target peptides in all four charge-states. The resolution was set to 120,000 at 200 m/z, 3×10^6^ AGC target, and 50 ms maximum injection time (IT). Second, for targeted PRM acquisition, SIL peptides were analyzed using the RT info (±1.5 min) from the prior DDA experiments to schedule the precursor list. MS1 data was recorded over the mass range from 150 to 1,450 m/z with a resolution of 60,000 at 200 m/z, with 3×10^6^ AGC target and 25 ms maximum IT. MS2 data was acquired using a resolution of 60,000 at 200 m/z, a normalized AGC target of 1000% (or 1×10^6^), and the maximum injection time mode was set to dynamic. The dynamic RT feature using the PRTC mixture and real-time lock mass calibration were enabled. Finally, an assay-external reference library was generated from PRM-based MS analysis of heavy synthetic peptides. The resulting experimental spectral library was used to confirm peptide identity in *Skyline* based on the NSA (experimental NSA ≥ 0.7; transitions ≥ 3) and the RT obtained from SIL peptides. Of note, spike-in of SIL peptides into the biological samples (plasma or fresh-frozen tumor tissue) was not feasible due to insufficient remaining material. Finally, variants resulting in neoepitope candidates were manually inspected in Integrative Genomics Viewer (version 2.19.7) with a mapping quality threshold of 1 and filtering out duplicated reads.

### Identification of tumor-associated antigens

Wild-type peptides returned by Spectronaut from all healthy and diseased individuals were queried against the HLA Ligand atlas^66^ and a database from IEDB^69^ containing linear peptides of human source confirmed in HLA I ligand assays. Peptides not found in these databases were further filtered based on the predicted HLA-binding affinity to retain only predicted HLA-binders (EL rank ≤ 2%). HLA ligands were manually evaluated in Skyline based on their *in silico* NSA (≥0.8) and transition counts (≥3). The source genes of HLA ligands were then queried against a curated TAA reference list. The reference TAA list was curated from the cancer tumor database (CTdatabase)^70^ (only “testis-restricted” TAAs) and Tumor T cell Antigen database (TANTIGEN)^71^ (only “Differentiation”, “Overexpressed”, “Shared tumor-specific” TAAs), for a total set of 163 unique TAAs. Among the 163 TAAs, 47 CTAs were classified according to Shraibman et al.^17^, who classified CTAs as genes with germline-restricted expression showing low expression in normal tissues (<9 GCRMA units) and significantly elevated expression in tumor tissues based on BioGPS data.

### Immunogenicity testing

To assess the immunogenic potential of neoepitopes identified by immunopeptidomics, interferon-γ (IFN-γ) enzyme-linked immunospot (ELISpot) and intracellular cytokine staining assays were performed as described previously^64^.

In brief, expanded PBMCs from HLA-(supertype-)matched healthy platelet donors (n = 7) of the Hannover Medical School (MHH) Institute of Transfusion Medicine and Transplant Engineering of Hannover Medical School (Ethical vote number: 3639-2017) were stimulated with synthetic neoepitopes of >95% purity and at a final concentration of 10 μg mL^−1^ (Research Group GMP & T Cell Therapy, DKFZ). On day 12, 1-2 × 10^5^ cells per well were seeded on ELISpot plates coated with 1:500 anti-human IFN-γ (clone 1-D1K; Mabtech) and restimulated with the respective peptide. Stimulation with concanavalin A (Sigma Aldrich, 2 μg/mL) and DMSO (Sigma Aldrich, 1 μL/mL) were used as positive and negative controls, respectively. After 24 h, 1:1000 biotinylated anti-human IFN-γ (clone 7 B6-1-Biotin; Mabtech), 1:2000 Streptavidin-Alkaline Phosphatase solution (Mabtech), and substrate (NBT/BCIP; Millipore) was used to develop the ELISpot. The number of spots was analyzed with an automated ImmunoSpot reader (CTL-Immunospot S6 Ultra-UV). The number of spot-forming units (SFU) per 1 × 10^6^ cells was calculated for each well and the SFU fold-change in peptide-stimulated wells relative to the background control was determined (stimulation index, SI). A peptide-specific T-cell response was considered positive when SI>3 and SFU>200 spots per 1×10^6^ cells.

T cell lines with positive ELISpot responses were split, and re-stimulated with 10 µg/mL of the respective peptide or mock-stimulated with 0.1% DMSO. GolgiStop (1:10, BD) and GolgiPlug (1:15, BD) were added after stimulation, and cells were incubated for 5 h. After surface staining for CD3 (OKT3, Biolegend), CD4 (RPA-T4, Biolegend), and CD8 (RPA-T8, BD) and viability NIR dye (Thermo Fisher), cells were fixed, permeabilized, and stained for intracellular IFN-γ (4S.B3, Biolegend) and TNFα (Mab11, BD). Samples were acquired on a BD FACS Canto™ II and analyzed using FlowJo v10.4.

## Acknowledgement

Rebecca Köhler is thanked for excellent technical support. We also want to acknowledge the excellent services and support provided by the DKFZ Proteomics Core Facility. Figures were created in BioRender (Kienzle, A., 2026; https://BioRender.com/qt8p4mo).

