## Supplementary Material for "Mutanome-guided immunopeptidomics of blood plasma for neoepitope detection in solid tumors is constrained by cfDNA variant calling sensitivity and MS detection limits"

### Appendix

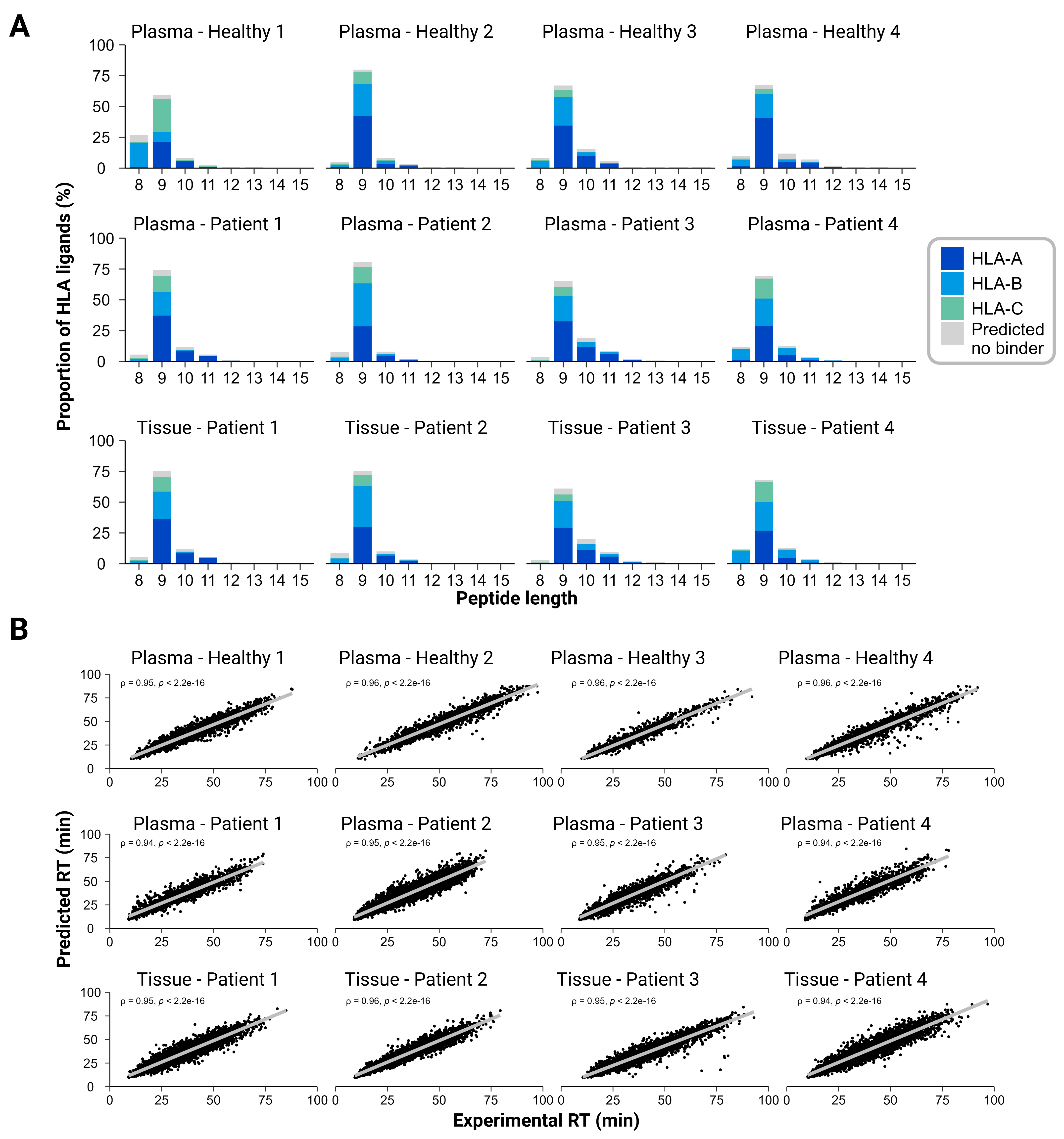

**Supplementary Figure 1: Immunopeptidomics of wild-type peptides**

**A** Length distribution and relative proportions of best-fit HLA alleles of wild-type HLA ligands from immunopeptidomic analysis of plasma from healthy individuals (n = 4) as well as tissue and plasma of patients (n = 4). HLA-binding affinity, scored by EL ranks (threshold: ≤ 2%), was predicted by NetMHCpan4.1.

**B** Correlation, based on the Spearman rank test, of predicted retention times (RTs) from DeepLC vs. experimental RTs (Spectronaut) of peptides per healthy individual (plasma only) and patient (tissue or plasma).

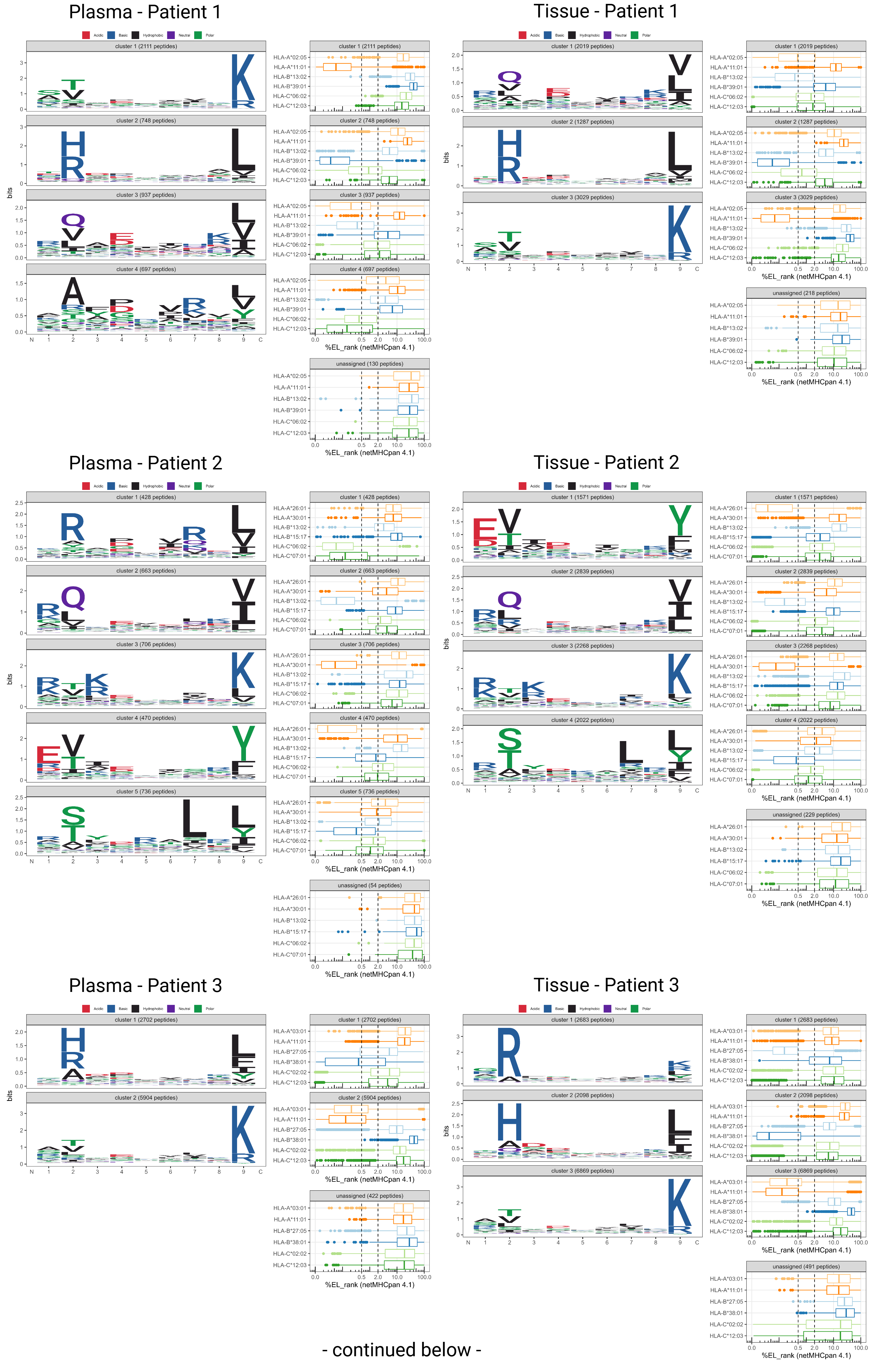

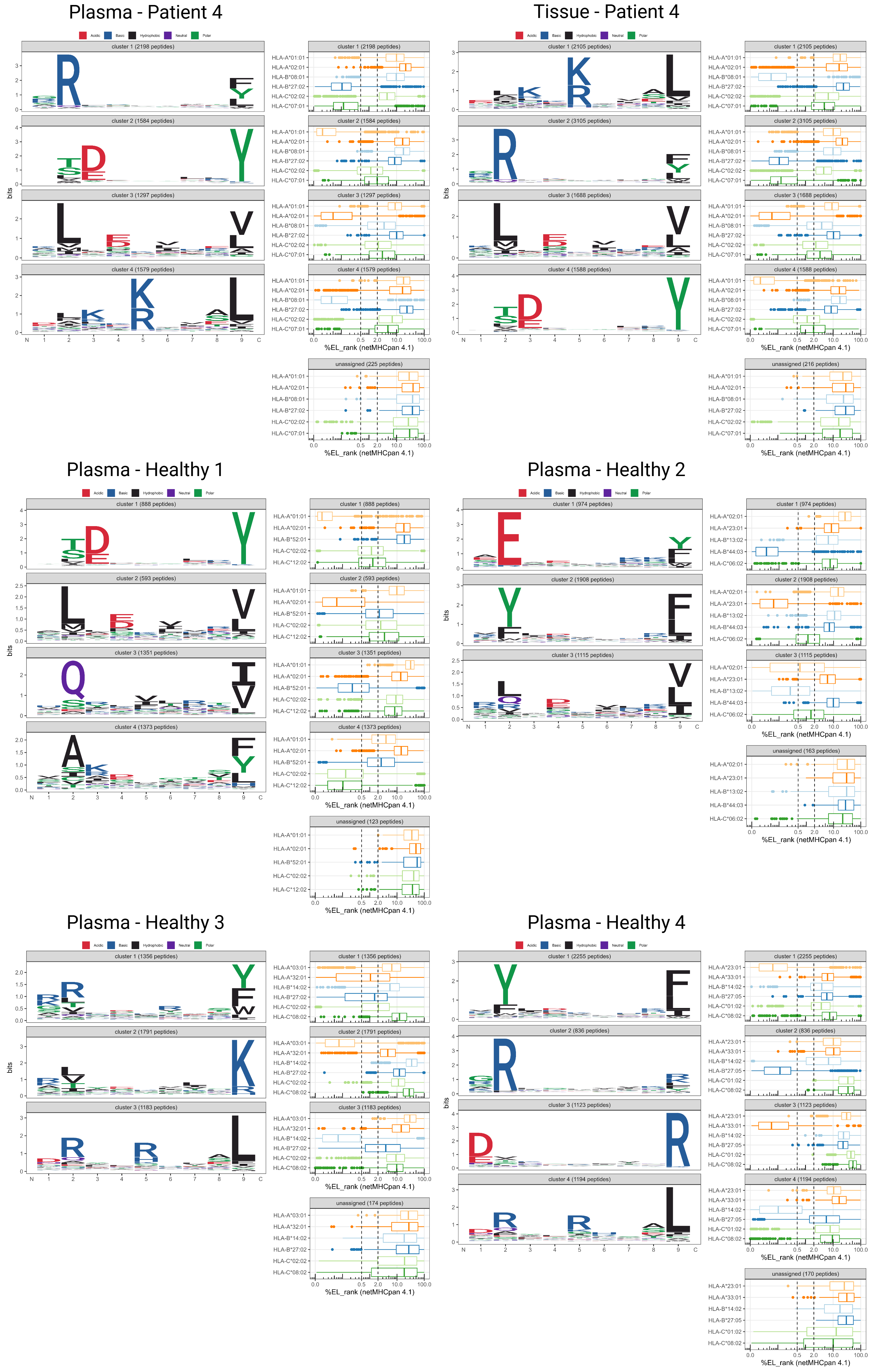

**Supplementary Figure 2: Gibbs clusters of wild-type peptides**

Gibbs sequence clustering of immunopeptidomics-identified wild-type HLA ligands from tissue and/or plasma of four patients and four healthy individuals. The clustering effectively segregates peptides into groups with shared sequence motifs that correlate with specific HLA binding preferences, reflecting the donor’s HLA type. For each cluster, box plots display the predicted HLA-binding affinities (%EL rank scores, netMHCpan 4.1) of the peptides to individual HLA alleles. Notably, the assignment of peptides in one cluster to multiple predicted HLA-alleles is expected due to the similarity of certain sequence motifs (for example, HLA-C*06:02 and HLA-B*39:01).

|  |  | **HLA-A** | | **HLA-B** | | **HLA-C** | |
| --- | --- | --- | --- | --- | --- | --- | --- |
| **Patients** | **1** | 02:05 | 11:01 | 13:02 | 39:01 | 06:02 | 12:03 |
|  | **2** | 26:01 | 30:01 | 13:02 | 15:17 | 06:02 | 07:01 |
|  | **3** | 03:01 | 11:01 | 27:05 | 38:01 | 02:02 | 12:03 |
|  | **4** | 01:01 | 02:01 | 08:01 | 27:02 | 02:02 | 07:01 |
| **Healthy** | **1** | 01:01 | 02:01 | 52:01 |  | 02:02 | 12:02 |
|  | **2** | 02:01 | 23:01 | 13:02 | 44:03 | 04:01 | 06:02 |
|  | **3** | 03:01 | 32:01 | 14:02 | 27:02 | 02:02 | 08:02 |
|  | **4** | 23:01 | 33:01 | 14:02 | 27:05 | 01:02 | 08:02 |

**Supplementary Table 1: HLA-genotypes**

Four-digit alleles of HLA-A, -B, and -C from four healthy individuals and four cancer patients, obtained from *NGS*-based HLA-genotyping.

|  |  | **MS detection source** | **Parental gene** | **Identified peptide** | **Caner testis antigen** | ***In silico***  **NSA** | **Transition count** | **RT (min)** | **Peptide charge state** | **Best-fit HLA allele** | **EL rank for best-fit HLA allele (%)** |
| --- | --- | --- | --- | --- | --- | --- | --- | --- | --- | --- | --- |
| **Patients** | 1 | Tissue | BIRC5 | CTPERMAEA | Yes | 0.87 | 7 | 13.6 | 2 | HLA-A02:05 | 1.705 |
|  |  | Blood | TP53 | CTYSPALNK | No | 0.82 | 5 | 27 | 2 | HLA-A11:01 | 0.017 |
|  |  | Tissue | TP53 | CTYSPALNK | No | 0.91 | 6 | 27.4 | 2 | HLA-A11:01 | 0.017 |
|  |  | Tissue | CSPG4 | QLYSGRLQV | No | 0.86 | 5 | 48.1 | 2 | HLA-B13:02 | 0.029 |
|  |  | Blood | CCND1 | RAYPDANLL | No | 0.87 | 5 | 48.9 | 2 | HLA-C12:03 | 0.045 |
|  |  | Tissue | CCND1 | RAYPDANLL | No | 0.8 | 4 | 48.9 | 2 | HLA-C12:03 | 0.045 |
|  |  | Blood | MUC2 | STDKQTCLK | No | 0.89 | 6 | 12.3 | 2 | HLA-A11:01 | 0.139 |
|  | 2 | Tissue | WDR46 | CRIDKSRKL | No | 0.87 | 9 | 14.5 | 3 | HLA-C06:02 | 0.016 |
|  |  | Tissue | MCL1 | ELYRQSLEI | No | 0.8 | 5 | 52.8 | 2 | HLA-B13:02 | 0.297 |
|  |  | Tissue | SFMBT1 | ESVMINGKY | No | 0.86 | 8 | 32.2 | 2 | HLA-A26:01 | 0.021 |
|  |  | Tissue | CA9 | FQSPVDIRP | No | 0.9 | 7 | 52.7 | 2 | HLA-B13:02 | 1.306 |
|  |  | Tissue | ERBB2 | FRNPHQALL | No | 0.85 | 9 | 34.8 | 2 | HLA-C06:02 | 0.001 |
|  |  | Blood | MUC2 | GQSCTAPKI | No | 0.87 | 3 | 26.3 | 2 | HLA-B13:02 | 0.048 |
|  |  | Tissue | MUC2 | GQSCTAPKI | No | 0.91 | 5 | 25.7 | 2 | HLA-B13:02 | 0.048 |
|  |  | Tissue | EPHA2 | GVRLPGHQK | No | 0.87 | 8 | 17 | 3 | HLA-A30:01 | 0.009 |
|  |  | Tissue | TOP2B | KFKAQTQLNK | No | 0.9 | 8 | 17.4 | 3 | HLA-A30:01 | 0.106 |
|  |  | Tissue | ZNF395 | KVLRSIVGI | No | 0.88 | 9 | 46.1 | 2 | HLA-B13:02 | 0.181 |
|  |  | Tissue | DDR1 | RAFQAMQV | No | 0.81 | 5 | 26.9 | 2 | HLA-B13:02 | 1.179 |
|  |  | Tissue | ADAM17 | RILKSPQEV | No | 0.88 | 8 | 26.8 | 2 | HLA-B13:02 | 0.03 |
|  |  | Tissue | CPSF1 | RQGELRISV | No | 0.92 | 5 | 39.3 | 2 | HLA-B13:02 | 0.003 |
|  |  | Tissue | CEACAM5 | RSDSVILNV | No | 0.89 | 7 | 49.8 | 2 | HLA-B13:02 | 0.304 |
|  |  | Tissue | ART4 | RTKDVHFNA | No | 0.82 | 6 | 18.5 | 3 | HLA-A30:01 | 0.007 |
|  | 3 | Blood | ABCC3 | AVVGPVGCGK | No | 0.85 | 7 | 29.6 | 2 | HLA-A03:01 | 0.194 |
|  |  | Tissue | ABCC3 | AVVGPVGCGK | No | 0.88 | 8 | 24.4 | 2 | HLA-A03:01 | 0.194 |
|  |  | Blood | TP53 | CTYSPALNK | No | 0.89 | 6 | 31.6 | 2 | HLA-A03:01 | 0.009 |
|  |  | Tissue | SCRN1 | GTPDPSRSIFK | No | 0.91 | 9 | 36.6 | 2 | HLA-A11:01 | 0.16 |
|  |  | Tissue | SART3 | QVISVTFEK | No | 0.83 | 4 | 47.9 | 2 | HLA-A11:01 | 0.009 |
|  |  | Blood | CCND1 | RAYPDANLL | No | 0.91 | 5 | 55.5 | 2 | HLA-C12:03 | 0.045 |
|  |  | Tissue | CCND1 | RAYPDANLL | No | 0.92 | 5 | 48.3 | 2 | HLA-C12:03 | 0.045 |
|  |  | Tissue | TOP2B | RLSYYGLRK | No | 0.92 | 6 | 27.9 | 3 | HLA-A03:01 | 0.024 |
|  |  | Blood | MRPL28 | RRAAIYDKY | No | 0.85 | 5 | 27.7 | 3 | HLA-B27:05 | 0.052 |
|  |  | Tissue | MRPL28 | RRAAIYDKY | No | 0.87 | 5 | 22.6 | 3 | HLA-B27:05 | 0.052 |
|  |  | Tissue | CYP1B1 | SHDDPEFREL | No | 0.81 | 7 | 38.4 | 2 | HLA-B38:01 | 0.012 |
|  |  | Blood | MUC2 | SVFSICHSK | No | 0.91 | 5 | 38.8 | 3 | HLA-A11:01 | 0.031 |
|  |  | Tissue | CPSF1 | TQEGVRITL | No | 0.81 | 8 | 49.8 | 2 | HLA-B38:01 | 0.211 |
|  |  | Blood | ENAH | YHHTGNNTF | No | 0.8 | 8 | 21.6 | 2 | HLA-B38:01 | 0.004 |
|  |  | Tissue | ENAH | YHHTGNNTF | No | 0.81 | 7 | 16.9 | 2 | HLA-B38:01 | 0.004 |
|  | 4 | Tissue | PRAME | ALLPSLSHC | Yes | 0.87 | 6 | 52.6 | 2 | HLA-A02:01 | 0.403 |
|  |  | Tissue | MLANA | ALMDKSLHV | No | 0.87 | 5 | 31.6 | 3 | HLA-A02:01 | 0.005 |
|  |  | Blood | CCNB1 | CSEYVKDIY | No | 0.88 | 8 | 43.8 | 2 | HLA-A01:01 | 0.105 |
|  |  | Tissue | CCNB1 | CSEYVKDIY | No | 0.86 | 8 | 41.5 | 2 | HLA-A01:01 | 0.105 |
|  |  | Tissue | MAGEA1 | EADPTGHSY | Yes | 0.9 | 5 | 18.9 | 2 | HLA-A01:01 | 0.005 |
|  |  | Tissue | TOP2A | ERVGLHKVF | No | 0.85 | 7 | 33.8 | 3 | HLA-C07:01 | 0.241 |
|  |  | Blood | MAGEA10 | GLYDGMEHL | Yes | 0.88 | 3 | 43.5 | 2 | HLA-A02:01 | 0.006 |
|  |  | Tissue | MAGEA10 | GLYDGMEHL | Yes | 0.85 | 4 | 41.2 | 2 | HLA-A02:01 | 0.006 |
|  |  | Blood | CYP1B1 | GRSMAFGHY | No | 0.8 | 5 | 26 | 3 | HLA-B27:02 | 0.229 |
|  |  | Tissue | CYP1B1 | GRSMAFGHY | No | 0.9 | 6 | 23.7 | 3 | HLA-B27:02 | 0.229 |
|  |  | Tissue | MAGEC2 | GVYAGREHFV | Yes | 0.8 | 7 | 33.8 | 2 | HLA-A02:01 | 1.401 |
|  |  | Blood | MAGEA4 | GVYDGREHTV | No | 0.91 | 6 | 24.6 | 2 | HLA-A02:01 | 0.34 |
|  |  | Tissue | MAGEA4 | GVYDGREHTV | No | 0.89 | 6 | 22.6 | 2 | HLA-A02:01 | 0.34 |
|  |  | Tissue | TYR | LLMEKEDYHSL | No | 0.86 | 9 | 38.5 | 2 | HLA-B08:01 | 0.236 |
|  |  | Tissue | CSPG4 | QLYSGRLQV | No | 0.86 | 5 | 48 | 2 | HLA-A02:01 | 0.076 |
|  |  | Blood | MRPL28 | RRAAIYDKY | No | 0.92 | 4 | 25.2 | 3 | HLA-B27:02 | 0.035 |
|  |  | Tissue | MRPL28 | RRAAIYDKY | No | 0.9 | 4 | 22.9 | 3 | HLA-B27:02 | 0.035 |
|  |  | Blood | TP53 | SDCTTIHY | No | 0.83 | 5 | 31.6 | 2 | HLA-A01:01 | 0.423 |
|  |  | Tissue | TP53 | SDCTTIHY | No | 0.81 | 5 | 29.7 | 2 | HLA-A01:01 | 0.423 |
|  |  | Tissue | DCT | SLDDYNHLV | No | 0.87 | 8 | 50.3 | 2 | HLA-A02:01 | 0.003 |
|  |  | Blood | ABCC3 | SRSPIYSHF | No | 0.8 | 4 | 40.2 | 3 | HLA-C07:01 | 0.006 |
|  |  | Tissue | ABCC3 | SRSPIYSHF | No | 0.85 | 9 | 37.3 | 2 | HLA-C07:01 | 0.006 |
|  |  | Blood | WDR46 | SRTGRHLAF | No | 0.83 | 3 | 26.2 | 3 | HLA-C07:01 | 0.022 |
|  |  | Tissue | WDR46 | SRTGRHLAF | No | 0.85 | 5 | 23.3 | 3 | HLA-C07:01 | 0.022 |
| **Healthy** | 1 | Blood | TOP2B | TAKEAKEYF | No | 0.89 | 10 | 30.9 | 2 | HLA-C02:02 | 0.128 |
|  | 2 | Blood | TOP2B | AESLKTLRY | No | 0.83 | 7 | 38.7 | 3 | HLA-B44:03 | 0.009 |
|  |  | Blood | MUC2 | GQSCTAPKI | No | 0.91 | 5 | 28.5 | 2 | HLA-B13:02 | 0.048 |
|  | 3 | Blood | TP53 | CTYSPALNK | No | 0.92 | 6 | 29 | 2 | HLA-A03:01 | 0.009 |
|  |  | Blood | TOP2B | RLSYYGLRK | No | 0.84 | 5 | 31.1 | 3 | HLA-A03:01 | 0.024 |
|  | 4 | Blood | CCND1 | DANLLNDRVLR | No | 0.88 | 6 | 51.8 | 3 | HLA-A33:01 | 0.332 |
|  |  | Blood | TOP2B | VALKVIHEL | No | 0.88 | 6 | 53.4 | 2 | HLA-C01:02 | 0.324 |

**Supplementary Table 2: Detection of tumor-associated HLA ligands in plasma and tissue**

Tumor-associated HLA ligands identified by immunopeptidomics in plasma and tissue samples from four cancer patients and four healthy individuals. TAA classification was based on a curated reference list from CTdatabase^67^ and TANTIGEN^68^. Cancer-testis antigens were defined according to Shraibman et al.^19^. HLA binding affinity was predicted by netMHCpan4.1.

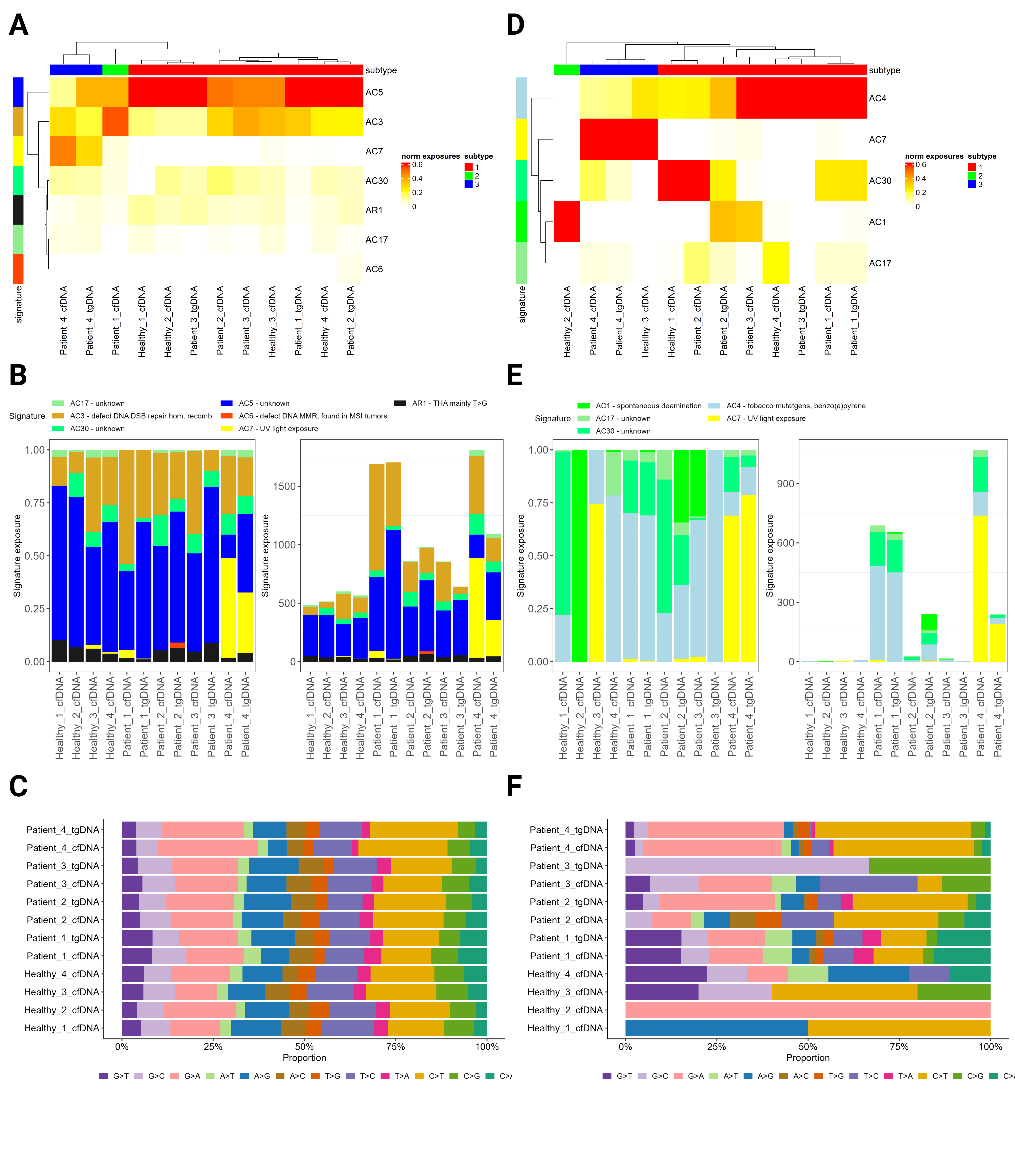
**Supplementary Figure 3: Mutational signatures**

**A-C** Mutational signatures of synonymous and non-synonymous variants from all variant calling pipelines, that passed the caller-specific built-in quality filters.

**D-F** Mutational signatures of synonymous and non-synonymous variants from all variant calling pipelines, that passed the manual filters using read coverage and strand bias filters.

**A,D** Hierarchical clustering (Manhattan distance, k = 3) of normalized COSMIC signature exposures (including artifacts) across samples (cfDNA, tgDNA) from healthy and diseased individuals. Color intensity reflects the relative contribution of each signature (AC: biological, e.g., aging; AR: artifacts, e.g., oxidation), as also shown in the legend of panel B and E.

**B,E** Stacked bar plot showing relative (left) and absolute (right) exposures to COSMIC signatures (including artifacts) per sample.

**C,F** Stacked bar plot of relative single-nucleotide variant types (reference to alternate allele change) across samples.

| **Patient** | **Neoepitope** | **MS search** | | | **Experimental validation** | | **Best predicted HLA-binding affinity** | **Variant calling** | | | | | |
| --- | --- | --- | --- | --- | --- | --- | --- | --- | --- | --- | --- | --- | --- |
|  |  | **Source** | **MS search engine** | **Search strategy** | **NSA** | **Transition count** |  | **Source** | **Variant caller** | **Read support (variant, reference) on tumor DNA** | **Read support (variant, reference) on leukocyte DNA** | **SOR for variant on tumor DNA** | **Parental gene**  **and mutation** |
| 1 | GSLQPLPPR | Plasma | Spectronaut | Library-free | 0.70 | 5 | HLA-A*11:01 (EL rank: 0.107%) | tgDNA | Sarek  M2 | 75,5 | 23,0 | 0.513 | chr12:109567094 (GRCh38) C🡪T  MMAB, ENSG00000139428, ENST00000545712, ENSP00000445920, p.Gly122Arg (missense) |
| 1 | GSLQPLPPR | Tissue | Spectronaut;  PEAKS | Library-free | 0.91 | 7 | HLA-A*11:01 (EL rank: 0.107%) | tgDNA | Sarek  M2 | 75,5 | 23,0 | 0.513 | chr12:109567094 (GRCh38) C🡪T  MMAB, ENSG00000139428, ENST00000545712  ENSP00000445920, p.Gly122Arg (missense) |
| 1 | SGAPPAPAAR | Plasma | PEAKS | Library-guided | 0.82 | 4 | HLA A*11:01 (EL rank: 1.747%) | tgDNA | Sarek  M2 | 538,14 | 59,0 | 2.775 | chr20: 20369331 (GRCh38) G🡪GGGCT  INSM1, ENSG00000173404, ENST00000310227  ENSP00000312631, p.Gly356AlafsTer199 (frameshift) |
| 1 | SGAPPAPAAR | Tissue | PEAKS | Library-free | 0.80 | 4 | HLA A*11:01 (EL rank: 1.747%) | tgDNA | Sarek  M2 | 538,14 | 59,0 | 2.775 | chr20: 20369331 (GRCh38) G🡪GGGCT  INSM1, ENSG00000173404, ENST00000310227  ENSP00000312631, p.Gly356AlafsTer199 (frameshift) |
| 2 | RTSLAALQL | Tissue | NeoDisc | Prediction | 0.91 | 4 | HLA-B*15:17 (EL rank: 0.108%) | tgDNA | NeoDisc  HC-M2-VS | 202,152 | 72,0 | 0.785 | chr2:43224585 (GRCh38) chr2: 43451724 (hg19) C🡪CG  ZFP36L2, ENSG00000152518.9, ENST00000282388.4  ENSP00000282388.3, p.407fs (frameshift) |
| 2 | RYKDLWTAY | Tissue | NeoDisc;  Spectronaut | Library-free | 0.91 | 6 | HLA-C*07:01 (EL rank: 0.161%) | tgDNA | NeoDisc HC-M1-M2-VS;  Sarek  M2 | NeoDisc:  82,60  Sarek:  90,66 | NeoDisc:  76,0  Sarek:  106,0 | NeoDisc:  0.638  Sarek:  0.680 | chr17:28690151 (GRCh38) chr17: 27017169 (hg19) C🡪T  SUPT6H, ENSG00000109111.16, ENST00000347486.8  ENSP00000338143.4, p.Arg1138Trp (missense) |
| 3 | KVSSIFFINK | Tissue | NeoDisc | Library-free | 0.81 | 6 | HLA-A*03:01 (EL rank: 0.079%) | tgDNA | NeoDisc  HC | 39,8 | 37,2 | 1.044 | chr2:9849441 (GRCh38) chr2: 9989570 (hg19)  TA🡪T TAF1B, ENSG00000115750.18, ENST00000263663.10, ENSP00000263663.4, p.62fs (frameshift) |

**Supplementary Table 3: Overview of experimentally validated neoepitopes**

Neoepitopes, detected in DIA-MS data and confirmed by assay-external SIL peptide spectral matching, from (matched) plasma and tissue samples. HLA-binding was predicted with NetMHCpan-4.1; for each peptide, the allele with the lowest predicted EL rank is listed. Read support for variant and reference position on either tumor DNA or leukocyte DNA was obtained from the variant caller output. Abbreviations for variant callers: M1, Mutect1; M2, Mutect2; VS, VarScan2; HC, HaplotypeCaller. StrandOddsRatio (SOR) was calculated using GATK VariantAnnotator from the sample BAM file.

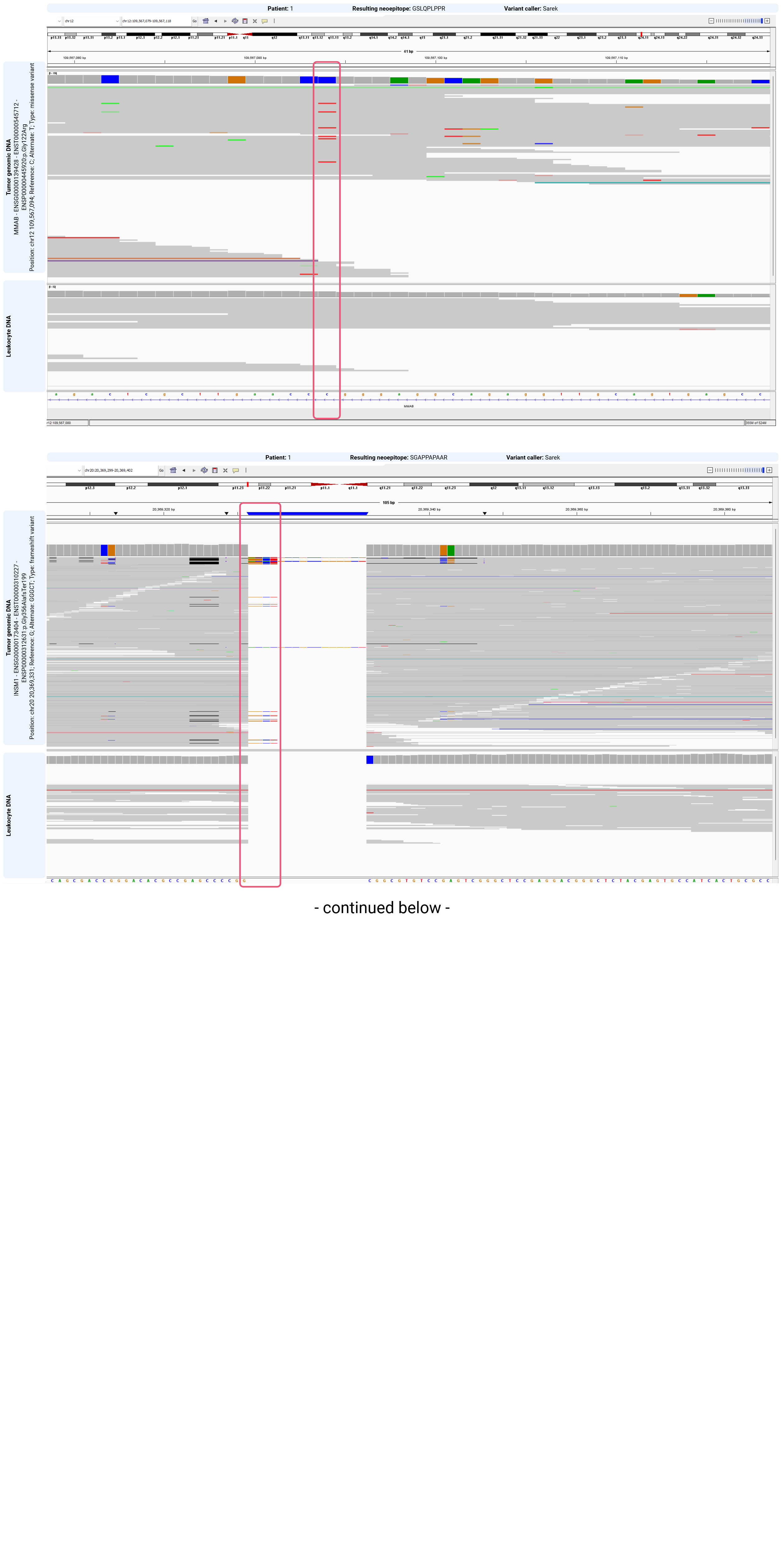

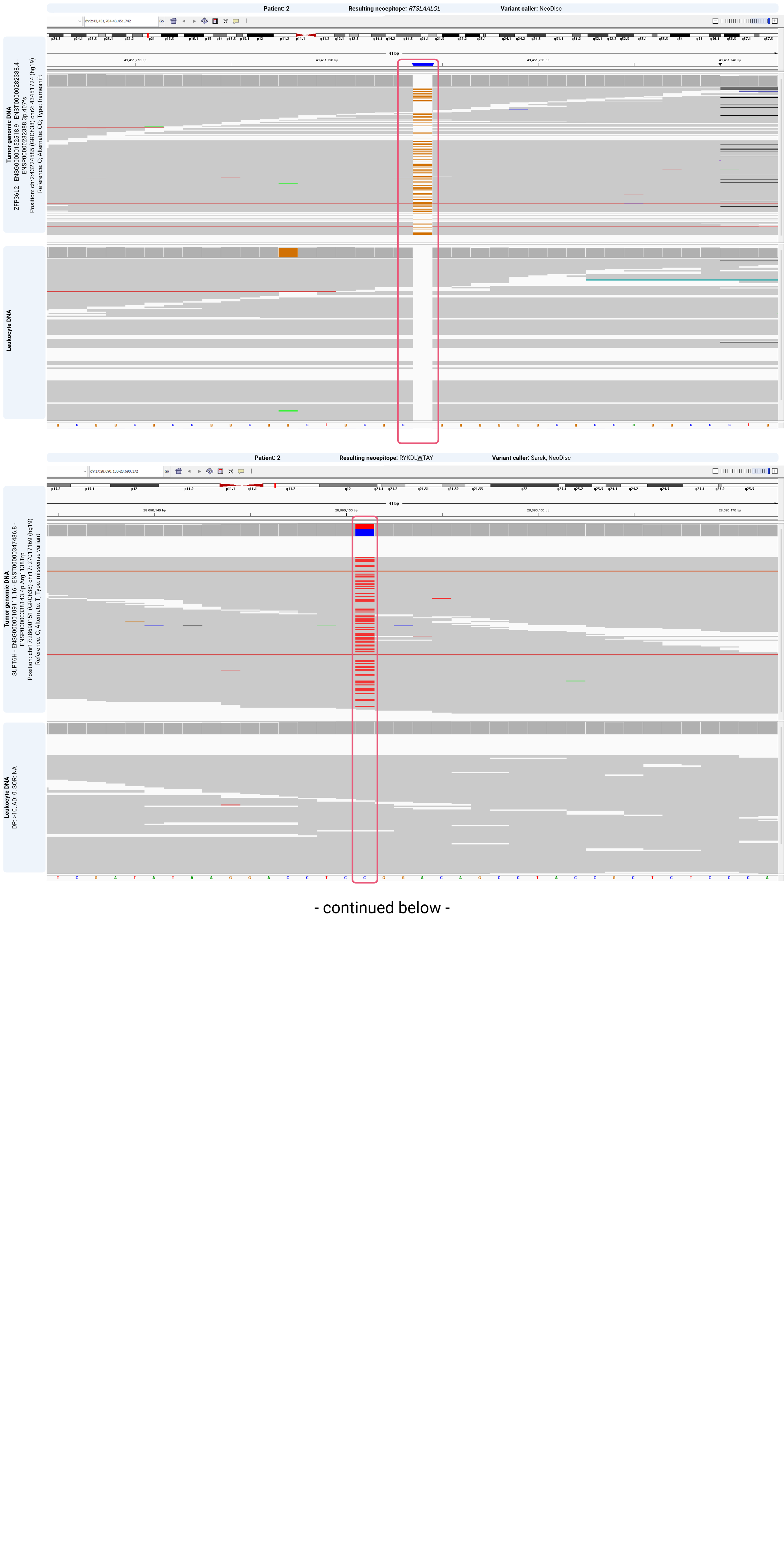

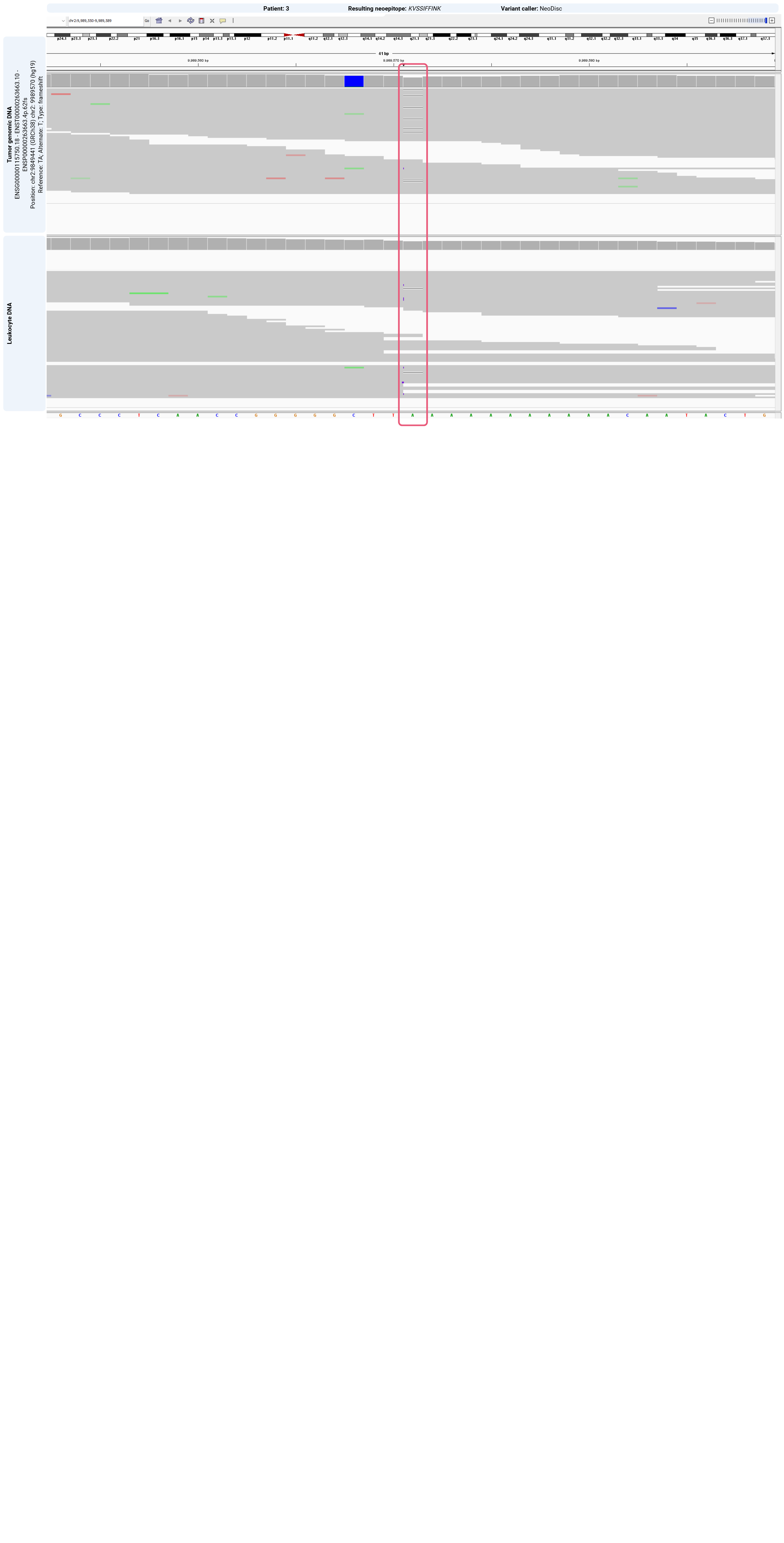

**Supplementary Figure 4: Manual evaluation of variants leading to neoepitope candidates**

Manual inspection of variant positions underlying neoepitope candidates on the DNA level in the *Integrative Genomics Viewer*. For each candidate, read support for the mutation of tumor DNA (cfDNA or tgDNA) and matched normal (leukocyte) DNA is shown. Variant positions from NeoDisc were based on the hg19 genome, while positions from Sarek were based on GRCh38.

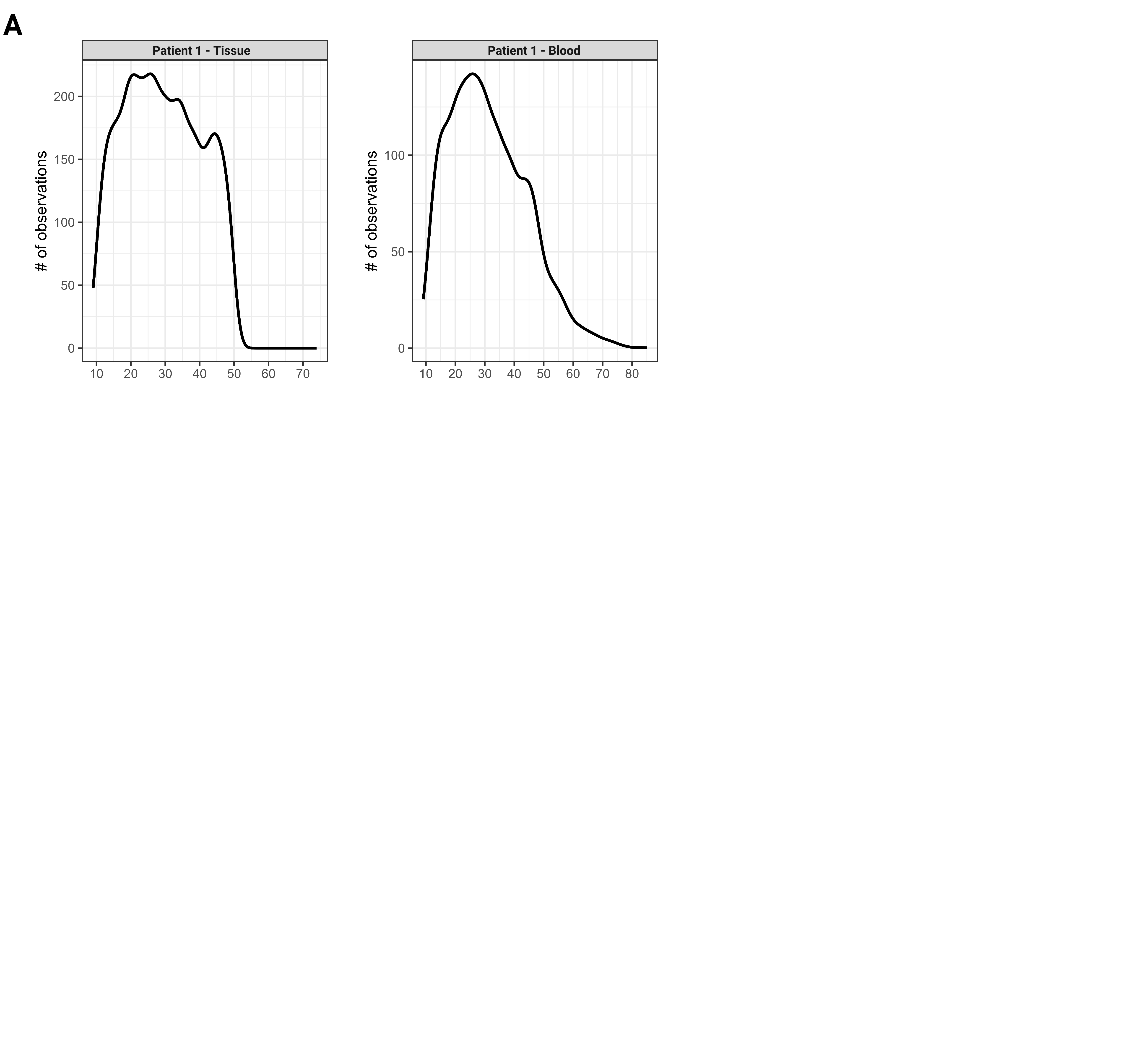

**Supplementary Figure 5: Retention time distribution of peptides**

Number of observations, corresponding to wild-type peptide IDs, versus retention time, in the plasma and tissue sample of patient 1.

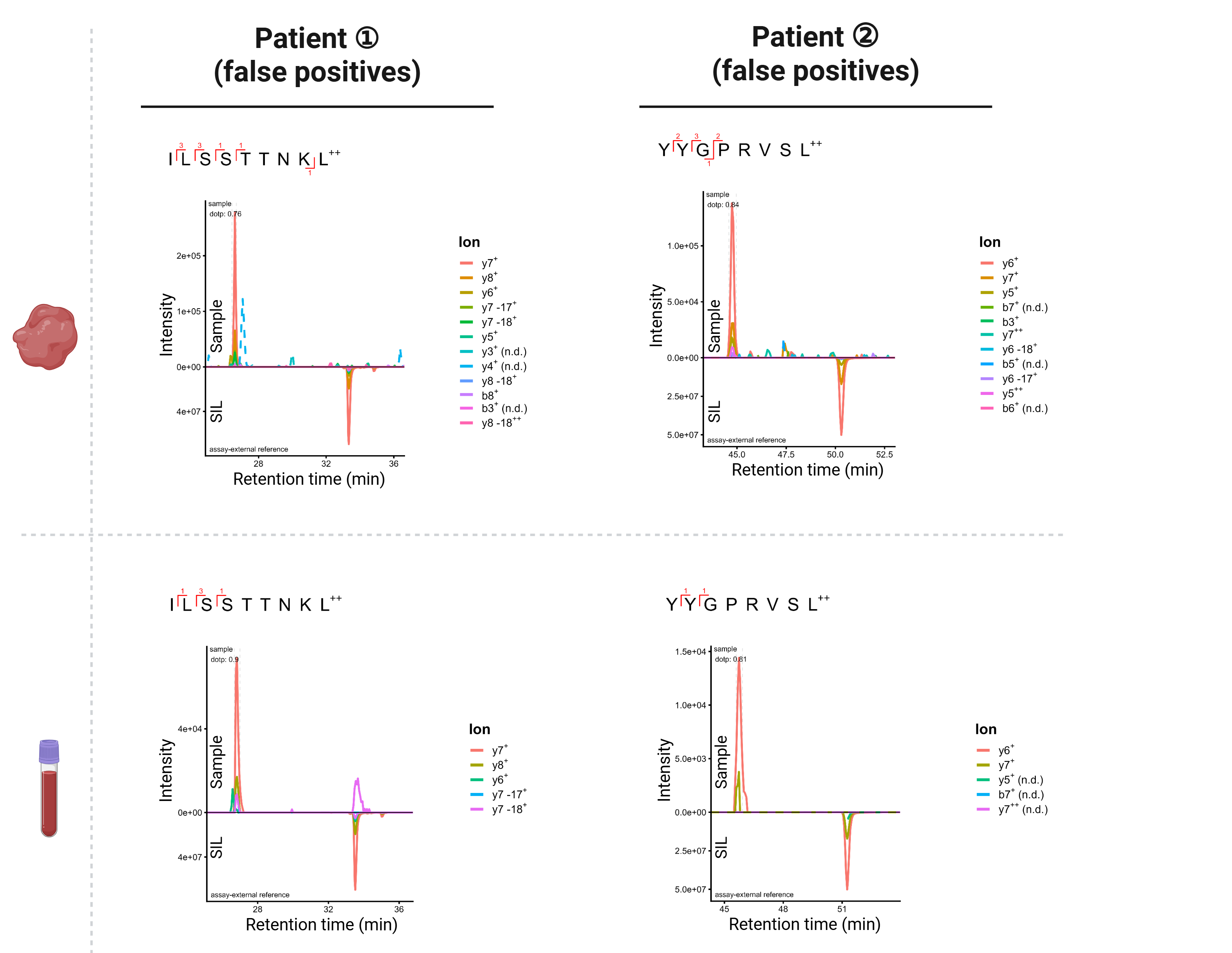

**Supplementary Figure 6: False-positive neoepitope hits**

Neoepitope candidates from DIA-MS data (top) classified as false-positives after validation using SIL peptides (bottom).

| **Patient** | **Neoepitope** | **Predicted HLA-binding affinity** | **HLA-genotypes of HLA-(supertype-)matched healthy donors** |
| --- | --- | --- | --- |
| 1 | GSLQPLPPR | HLA-A*11:01 (EL rank: 0.107%) | D6: HLA-A*11:01, HLA-A*68:01, HLA-B*41:01, HLA-B*44:02, HLA-C*07:04, HLA-C*17:01  **D11: HLA-A*11:01, HLA-A*23:01, HLA-B*18:03, HLA-B*44:03, HLA-C*04:01, HLA-C*07:01**  D16: HLA-A*02:01, HLA-A*11:01, HLA-B*27:05, HLA-B*35:03, HLA-C*02:02, HLA-C*04:01  D25: HLA-A*03:01, HLA-A*11:01, HLA-B*40:01, HLA-B*50:01, HLA-C*03:04, HLA-C*06:02 |
| 1 | SGAPPAPAAR | HLA A*11:01 (EL rank: 1.747%) | D6: HLA-A*11:01, HLA-A*68:01, HLA-B*41:01, HLA-B*44:02, HLA-C*07:04, HLA-C*17:01  **D11: HLA-A*11:01, HLA-A*23:01, HLA-B*18:03, HLA-B*44:03, HLA-C*04:01, HLA-C*07:01**  D16: HLA-A*02:01, HLA-A*11:01, HLA-B*27:05, HLA-B*35:03, HLA-C*02:02, HLA-C*04:01  D25: HLA-A*03:01, HLA-A*11:01, HLA-B*40:01, HLA-B*50:01, HLA-C*03:04, HLA-C*06:02 |
| 2 | RTSLAALQL | HLA-A*30:01 (EL rank: 0.598%)  HLA-B*13:02 (EL rank: 1.085%)  HLA-B*15:17 (EL rank: 0.108%)  HLA-C*07:01 (EL rank: 1.564%) | D5: HLA-A*01:01, HLA-A*26:01, HLA-B*08:01, HLA-B*08:01, HLA-C*07:01, HLA-C*07:01  D8: HLA-A*01:01, HLA-A*24:02, HLA-B*18:01, HLA-B*58:01, HLA-C*05:01, HLA-C*07:01  **D11: HLA-A*11:01, HLA-A*23:01, HLA-B*18:03, HLA-B*44:03, HLA-C*04:01, HLA-C*07:01**  D19: HLA-A*03:01, HLA-A*24:02, HLA-B*07:02, HLA-B*13:02, HLA-C*06:02, HLA-C*07:02 |
| 2 | RYKDLWTAY | HLA-A*26:01 (EL rank: 1.900%)  HLA-A*30:01 (EL rank: 0.198%)  HLA-C*07:01 (EL rank: 0.161%)  HLA-C*06:02 (EL rank: 0.504%) | D5: HLA-A*01:01, HLA-A*26:01, HLA-B*08:01, HLA-B*08:01, HLA-C*07:01, HLA-C*07:01  D8: HLA-A*01:01, HLA-A*24:02, HLA-B*18:01, HLA-B*58:01, HLA-C*05:01, HLA-C*07:01  **D11: HLA-A*11:01, HLA-A*23:01, HLA-B*18:03, HLA-B*44:03, HLA-C*04:01, HLA-C*07:01**  D19: HLA-A*03:01, HLA-A*24:02, HLA-B*07:02, HLA-B*13:02, HLA-C*06:02, HLA-C*07:02 |
| 3 | KVSSIFFINK | HLA-A*03:01 (EL rank: 0.079%)  HLA-A*11:01 (EL rank: 0.132%) | D6: HLA-A*11:01, HLA-A*68:01, HLA-B*41:01, HLA-B*44:02, HLA-C*07:04, HLA-C*17:01  **D11: HLA-A*11:01, HLA-A*23:01, HLA-B*18:03, HLA-B*44:03, HLA-C*04:01, HLA-C*07:01**  D16: HLA-A*02:01, HLA-A*11:01, HLA-B*27:05, HLA-B*35:03, HLA-C*02:02, HLA-C*04:01  D19: HLA-A*03:01, HLA-A*24:02, HLA-B*07:02, HLA-B*13:02, HLA-C*06:02, HLA-C*07:02  D25: HLA-A*03:01, HLA-A*11:01, HLA-B*40:01, HLA-B*50:01, HLA-C*03:04, HLA-C*06:02 |

**Supplementary Table 4: Immunogenicity assessment of neoepitopes.**

Neoepitopes identified from plasma and/or tissue (Supplementary Table 3) were tested for immunogenicity using PBMCs from seven HLA-(supertype-)matched healthy donors. HLA genotypes of donors tested per neoepitope are listed. Donor D11 (bold) showed reactivity against *KVSSIFFINK* and *RTSLAALQL*; the remaining six donors showed no reactivity against any tested neoepitope. HLA I binding affinities were predicted with NetMHCpan-4.1; only alleles with EL rank < 2% are shown. HIV-derived peptides binding to HLA-A03:01 (*RLRPGGKKK*) and HLA-A11:01 (*PLRPMTYK*) were included as negative controls (SI<3 and SFU<200 spots per 1×10⁶ cells). DMSO served as vehicle control.
