## Supplementary material for "Mutanome-guided immunopeptidomics of blood plasma for neoepitope detection in solid tumors is constrained by cfDNA variant calling sensitivity and MS detection limits": Highlights

- Mutation-derived neoepitopes were directly detected in patient blood plasma at the MS detection limit by immunopeptidomics using tumor tissue DNA mutanomes
- Detected neoepitopes from plasma were also identified in matching tumor tissue
- While there was a substantial overlap in wild-type HLA ligands between tissue and plasma, the overlap between tissue- and blood-borne mutanomes was limited
- Validation with stable isotope-labeled peptides is essential to eliminate false-positives and confirm true detection of mutation-derived neoepitopes
- At present, tumor tissue remains the most reliable source for variant calling and neoepitope detection
